# Cooperation between MYC and β-catenin in liver tumorigenesis requires Yap/Taz

**DOI:** 10.1101/819631

**Authors:** Andrea Bisso, Marco Filipuzzi, Gianni Paolo Gamarra Figueroa, Giulia Brumana, Francesca Biagioni, Mirko Doni, Giorgia Ceccotti, Nina Tanaskovic, Marco Jacopo Morelli, Vera Pendino, Fulvio Chiacchiera, Diego Pasini, Daniela Olivero, Stefano Campaner, Arianna Sabò, Bruno Amati

## Abstract

**Background & Aims:** Activation of *MYC* and *CTNNB1* (encoding β-catenin) can co-occur in liver cancer, but how these oncogenes cooperate in tumorigenesis remains unclear.

**Approach & Results:** We generated a mouse model allowing conditional activation of MYC and WNT/β-catenin signaling (through either β-catenin activation or Apc loss) upon expression of CRE recombinase in the liver, and monitored their effects on hepatocyte proliferation, apoptosis, gene expression profiles and tumorigenesis. Conditional activation of WNT/β-catenin signaling strongly accelerated MYC-driven carcinogenesis in the mouse liver. Both pathways also cooperated in promoting cellular transformation *in vitro*, demonstrating their cell-autonomous action. Short-term induction of MYC and β-catenin in hepatocytes followed by RNA-seq profiling allowed the identification of a “Myc/β-catenin signature”, composed of a discrete set of Myc-activated genes whose expression increased in presence of active β-catenin. Notably this signature enriched for targets of Yap and Taz, two transcriptional co-activators known to be activated by WNT/β-catenin signaling, and to cooperate with MYC in mitogenic activation and liver transformation. Consistent with these regulatory connections, Yap/Taz accumulated upon Myc/β-catenin activation and were required not only for the ensuing proliferative response, but also for tumor cell growth and survival. Finally, the Myc/β-catenin signature was enriched in a subset of human hepatocellular carcinomas characterized by comparatively poor prognosis.

**Conclusions:** Yap and Taz mediate the cooperative action of Myc and β-catenin in liver tumorigenesis. This warrants efforts toward therapeutic targeting of Yap/Taz in aggressive liver tumors marked by elevated Myc/β-catenin activity.

## Introduction

Liver cancers, among which hepatocellular carcinoma (HCC) is the predominant form, are the second cause of cancer-related mortality worldwide (1). Heterogeneity in genomic alterations in HCCs, as revealed by recent large-scale genome sequencing efforts, renders the understanding of driving molecular events and the design of targeted therapeutic strategies very challenging, with HCC patients still having limited therapeutic options (1, 2).

One of the most frequently altered pathways in HCC is WNT/β-catenin signaling (1–3). In this pathway, also known as “canonical” WNT signaling, the exposure of cells to WNT ligands leads to stabilization of the transcriptional coactivator β-catenin, which is then free to translocate to the nucleus and activate target genes in association with DNA-binding factors of the TCF family (4). β-catenin turnover is regulated by the so-called “destruction complex”, a cytoplasmic assembly that includes - among others - Axin1, APC, the ubiquitin ligase β-TrCP and the kinases CK1*α*/*δ* and GSK3*α*/β. In the absence of Wnt ligands, these kinases phosphorylate β-catenin at several sites, creating a discrete phospho-degron motif that acts as a docking site for β-TrCP, thus triggering ubiquitination and proteasome-mediated degradation of β-catenin (4). Activation of the WNT/β-catenin pathway in cancer can occur either via enhanced exposure to WNT ligands, or through genetic lesions in its core components: the latter include loss of APC, most frequent in colorectal cancer (4), or activating mutations in the β-catenin gene *CTNNB1*, as commonly observed in HCC (1–3).

The *MYC* proto-oncogene is one of the transcriptional targets of β-catenin/TCF, and functions as a key downstream effector of WNT/β-catenin signaling in several tissues, such as the small intestine, T cells and lung (5–8). In the liver, however, this epistatic relationship may not hold true: in particular, APC loss and the consequent activation of β-catenin do not induce *Myc* expression (9) and *Myc* deletion does not suppress the effects of APC loss on either hepatocyte proliferation (10) or liver zonation (11). Moreover, there are indications that the two pathways can be independently activated and cooperate in tumorigenesis (12). In particular, *MYC* amplification and *CTNNB1* mutations showed a tendency toward co-occurrence in either adult HCC or aggressive childhood hepatoblastoma (3, 5, 13), and MYC-driven HCC in experimental mouse models frequently acquired activating mutations in *Ctnnb1* (5, 14, 15).

In order to unravel the functional cross-talk between MYC and WNT/β-catenin signaling in liver tumorigenesis, we generated a new mouse model allowing conditional activation of MYC and β-catenin in hepatocytes. This model demonstrated a strong cooperativity between the two oncogenes in promoting liver tumorigenesis. Our data indicate that this cooperation occurs mainly through unrestrained proliferation of liver cells, and requires activation of the transcriptional co-factors Yap and Taz.

## Materials and Methods

### Mice

Alb-CreER^T2^ mice (termed SA-Cre-ERT2 in the original publication (16)) were a kind gift from Pierre Chambon, Ctnnb1*^Δ^*^Ex3^ mice were a kind gift from Makoto Taketo (17), Taz^f/f^ mice (18) were a kind gift from Stefano Piccolo, Apc^f/f^ (19) mice were a kind gift of Eduard Battle, R26-lsl-CAG-MYC-ires-hCD2* (20) mice were purchased from Jackson Laboratory (Stock No: 020458), R26-lsl-EYFP mice were purchased from Jackson Laboratory (Stock No: 006148) and backcrossed into the C57BL/6 background, and Yap^f/f^ mice were purchased from the KOMP Knockout mouse project (https://www.komp.org). For tumor-free survival analysis Alb-CreER^T2^;β-cat^Ex3^;R26-lslMYC mice were monitored 3 times per week and sacrificed when showing abdominal enlargement, indicative of hepatomegaly due to tumor formation. The same procedure was applied to β-cat^Ex3^;R26-lslMYC mice injected at 6-8 weeks of age by tail vein injection with low-titer (10^9^) AAV8-TBG-CRE particles (University of Pennsylvania Vector Core, #AV-8-PV1091). For short-term liver-specific activation of the various alleles, high-titer (10^11^) AAV8-TBG-CRE particles were given to 6- to 8-week-old mice, and the mice sacrificed after 2, 4, or 8 days, as specified in the text or figures. Tumor nodules or liver parenchyma were dissected and either processed freshly, or frozen and stored at −80°C until further analysis. CD1-nude mice (purchased from Charles River Laboratories) were injected subcutaneously with 3*10^6^ 3T9^MycER;S33Y^ cells, and provided with food containing 400 ppm/kg Tamoxifen (Envigo, #TD.55125.I) and/or drinking water with 2 mg/ml doxycyline hydrate (Sigma-Aldrich, #D9891-100G), as indicated in the text. Mice were monitored 3 times per week and sacrificed when showing tumor masses of 1 cm^3^. Experiments involving animals were done in accordance with the Italian Laws (D. lgs. 26/2014), which enforces Dir. 2010/63/EU (Directive 2010/63/EU of the European Parliament and of the Council of 22 September 2010 on the protection of animals used for scientific purposes) and authorized by the Italian Minister of Health with projects 81/2016-PR and 726/2017-PR.

Additional Methods can be found in Supplementary Material and Methods.

## Results

### MYC and β-catenin cooperate in liver tumorigenesis

In line with previous observations (21) analysis of the TCGA database showed that *MYC* amplification and *CTNNB1* mutations co occur in a fraction of human HCCs (Supplementary Figure 1A) (2, 3). To model this scenario, we took advantage of the CRE-activated alleles R26-lsl-CAG-MYC-ires-hCD2* (hereafter R26-lslMYC) (20) and Ctnnb1*^Δ^*^Ex3^ (hereafter *β-cat*^Ex3^) (17) (Supplementary Figure 1B). Recombination of the *β-cat*^Ex3^ allele leads to loss of residues 14-89, spanning the phospho-degron of mouse β-catenin, thus mimicking the cancer-associated mutations observed in the human gene. To circumvent the known lethal effects of β-catenin activation in the whole liver (22), we sought to obtain a sporadic activation of our alleles in a minority of the hepatocytes, which was achieved by two complementary means: (i.) exploiting the basal leakiness of the CreER^T2^ fusion protein in the absence of its pharmacological activator, and (ii.) infection with low titer CRE-expressing Adeno-Associated viral (AAV) particles.

The Alb-CreER^T2^ transgene drives constitutive expression of the CreER^T2^ fusion protein in the liver, allowing controlled post-translational activation of the recombinase by exposure of the animals to Tamoxifen (TAM) (16). CreER^T2^ can occasionally show leakiness (23), causing sporadic recombination events: indeed, even if never exposed to TAM, compound Alb-CreER^T2^;R26-lslMYC mice developed highly penetrant, gender-independent multi-nodal liver tumors, with a median survival of 6 months (Figure 1A, Supplementary Figure 2A-B). Activation of the *β-cat*^Ex3^ allele had no effect on its own, as previously reported (22), but significantly accelerated tumorigenesis when combined with R26-lslMYC (Figure 1A; supplementary Figure 2A). As expected, the resulting tumors expressed MYC at high levels, the associated hCD2 reporter, as well as the shorter form of β-catenin encoded by the recombined *β-cat*^Ex3^ allele (Figure 1B; supplementary Figure 1B, 2C). We further assessed the expression of known MYC- and β-catenin-activated genes, confirming the activation of the two oncoproteins (Supplementary Figure 2D, E). The tumors also showed reduced expression of markers typically associated with differentiated hepatocytes (e. g. Ck-18, Albumin, and Hnf-4a) and increased expression of fetal liver markers (e. g. Afp; Supplementary Figure 2F). Finally, pathological analysis revealed no major differences between MYC-only and MYC/β-catenin tumors, both resembling human HCC with a trabecular pattern (Figure 1C; see Supplementary Table 1 for a detailed pathological description). We will thus generally refer to these tumors as mouse hepatocellular carcinomas (mHCCs).

**Figure 1.**
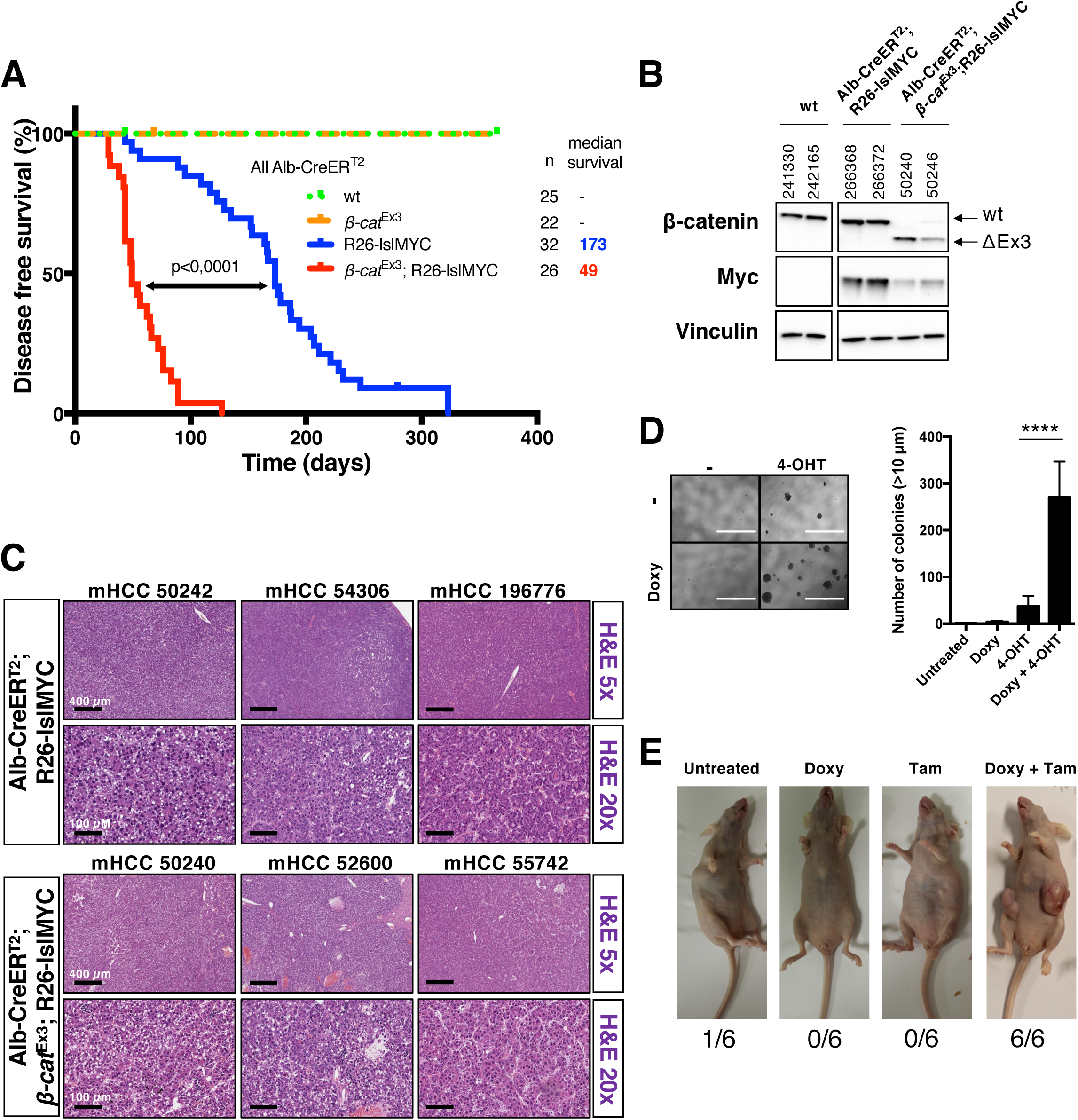
MYC and β-cat^Ex3^ activation cooperate in transformation in liver and fibroblasts. **(A)** Kaplan Meyer disease-free survival curves for mice of the indicated genotypes (all in the presence of the Alb-CreER^T2^ transgene). The number of mice (n) and the median survival are indicated. p-values were calculated with the log-rank test. **(B)** Western blot analysis of β-catenin and MYC protein expression in wild-type (wt) liver and representative R26-lslMYC and *β-cat*^Ex3^;R26-lslMYC liver tumor samples. Vinculin was used as loading control. Each tumor is identified by its unique reference number. **(C)** Hematoxylin and Eosin staining of representative liver sections from the indicated genotypes. Bars: 400 µm (H&E 5x) or 100 µm (H&E 5x). **(D)** Left: representative pictures of the colony forming assay of 3T9^MycER;S33Y^ cells plated in 50% (v/v) methylcellulose and cultured for 7-10 days in the presence of 1 μM Doxycycline and/or 400 nM 4-OHT, as indicated. This experiment was performed four times, each with technical triplicates. Bar: 1000 µm. Right: quantification of the total number of colonies with a diameter >10 μM in each condition. Bar plot represent the average and standard deviation for the 4 biological replicates. **(E)** Representative pictures of CD1-nude mice 6 weeks after injection of 3×10^6^ 3T9^MycER;S33Y^ cells and fed with Tamoxifen (Tam) and/or Doxycyline (Doxy), as indicated. The fraction of tumor-bearing mice in each experimental group is indicated below each photograph.

As a second approach to jointly activate MYC and β-catenin in hepatocytes, *β-cat*^Ex3^;R26-lslMYC and control mice were injected with recombinant AAV8-TBG-CRE particles, which show a high tropism for liver cells and express CRE recombinase under control of the hepatocyte-specific TBG promoter (24). This system can be used to achieve either sporadic (*∼*5-10% of the cells) or almost complete (>90%) infection of hepatocytes, as measured by CRE-induced activation of the R26-lsl-EYFP reporter allele 4 days after injection of 10^9^ (low titer) or 10^11^ (high titer) particles (Supplementary Figure 3A). When infected with AAV8-TBG-CRE at low titer, *β-cat*^Ex3^;R26-lslMYC mice developed tumors that resembled those arising in Alb-CreER^T2^;*β-cat*^Ex3^;R26-lslMYC animals, albeit with longer latency (Supplementary Figure 3B, C; Supplementary Table1) – owing most likely to activation of the oncogenes at a later stage of liver development.

Finally, as an alternative means to activate the WNT/β-catenin pathway, we used a conditional knockout allele of *APC* (*19*), bred this to homozygosity (*Apc^f/f^*) in combination with Alb-CreER^T2^ and R26-lslMYC, and monitored the mice over time in the absence of TAM: as observed above with activation of *β-cat*^Ex3^, deletion of APC had no effect on its own, but accelerated tumorigenesis in the presence R26-lslMYC (Supplementary Figure 2G). Altogether, our data show that oncogenic activation of MYC and WNT/β-catenin signaling cooperate in liver tumorigenesis.

We next wondered whether the cooperation between MYC and β-catenin is peculiar to hepatocytes or may be recapitulated in other cells. To address this question, we infected 3T9^MycER^ fibroblasts, which constitutively express a MycER^T2^ chimaera (25), with a doxycycline-inducible lentiviral vector encoding a 6xMyc-tagged S33Y-mutant version of β-catenin resistant to GSK3β-induced phosphorylation and proteasomal degradation (hereafter β-cat^S33Y^) (26). Immunoblot and RT-qPCR analysis of 3T9^MycER;S33Y^ cells confirmed the induction of β-cat^S33Y^ by doxycycline, as well as an increase in MycER levels – most likely due to stabilization of the protein – upon 4-hydroxytamoxifen (4-OHT) treatment (Supplementary Figure 4A). Importantly, β-catenin- and MYC-activated genes were induced upon doxycycline and 4-OHT treatment, respectively, indicating that the two transcription factors were active (Supplementary Figure 4B, C). The induction of β- cat^S33Y^ reduced cell proliferation in adherent cultures, independently from MycER activation (Supplementary Figure 4D); most importantly, however, co-activation of MycER^T2^ and β-cat^S33Y^ allowed colony formation in methylcellulose, indicating that the cells had acquired the capacity to proliferate in suspension (Figure 1D). 3T9^MycER;S33Y^ cells were then implanted subcutaneously in CD1-nude mice: in this setting, combined treatment of the animals with Doxycycline and Tamoxifen, but neither alone, led to tumor formation (Figure 1E). Thus, co-activation of MYC and β-catenin supports the malignant transformation of immortalized mouse fibroblasts.

Altogether, the above results demonstrate that MYC and WNT/β-catenin signaling cooperate in cellular transformation. In particular, we showed that the aberrant activation of MYC in the mouse liver promoted the development of undifferentiated hepatocellular carcinomas, whose latency was strongly reduced upon either activation of β-catenin, or deletion of APC. Most importantly, MycER and β-cat^S33Y^ also cooperated in the transformation of fibroblasts *in vitro*. Thus, besides its recently reported impact on immune surveillance (21), β-catenin also cooperates with MYC through cell-autonomous mechanisms.

### Concomitant activation of Myc and β-catenin induces of a pro-proliferative, Yap/Taz-related, transcriptional signature

Considering the prevalence of secondary transcriptional responses in established Myc-driven mHCCs (27), we sought to profile the short-term effects of MYC and β-cat^Ex3^ activation in hepatocytes. In this regard, short-term activation of CreER^T2^ in young Alb-CreER^T2^;*β-cat*^Ex3^;R26-lslMYC mice did not provide a clean model, owing to the possible confounding effects of pre-existing tumoral lesions. In order to circumvent this caveat, we focused on the short-term activation of R26-lslMYC and/or β-cat^Ex3^ with injection of high-titer AAV8-TBG-CRE particles. RNA-seq profiles were established 4 days after infection, a time at which both MYC and β-cat^Ex3^ were expressed in hepatocytes, showing clear nuclear localization and transcriptional activity (Supplementary Figure 5A-E). Of note, activation of *β-cat*^Ex3^ did not increase *Myc* mRNA levels (Supplementary Figure 5C), indicating that *Myc* is not regulated by WNT/β-catenin in hepatocytes, as previously reported (9, 10).

Activation of β-cat^Ex3^ alone induced limited transcriptional changes relative to AAV-infected wild-type livers, with only 98 differentially expressed genes (DEGs), while MYC regulated almost 3000 genes, either up or down (Figure 2A). This MYC-driven transcriptional program was largely unchanged upon co-activation of β-cat^Ex3^ (Figure 2B, Supplementary Figure 6A). Gene ontology analysis showed that MYC, either alone or in combination with β-catenin, elicited not only MYC-, but also E2F-, mTOR- and WNT/β-catenin-associated gene signatures: the latter was mobilized by either β-cat^Ex3^ or MYC alone, but responded with higher statistical significance in presence of both oncogenes (Figure 2C, Supplementary Table 2). In line with these findings, direct comparison between MYC/β-cat^Ex3^ and MYC-overexpressing livers yielded only 37 DEGs (33 up and 4 down, Supplementary Figure 6A), comprising canonical WNT/β-catenin targets (Axin2, Lgr5, Notum, Sp5, Tbx3 and Tcf7; Supplementary Table 2).

**Figure 2.**
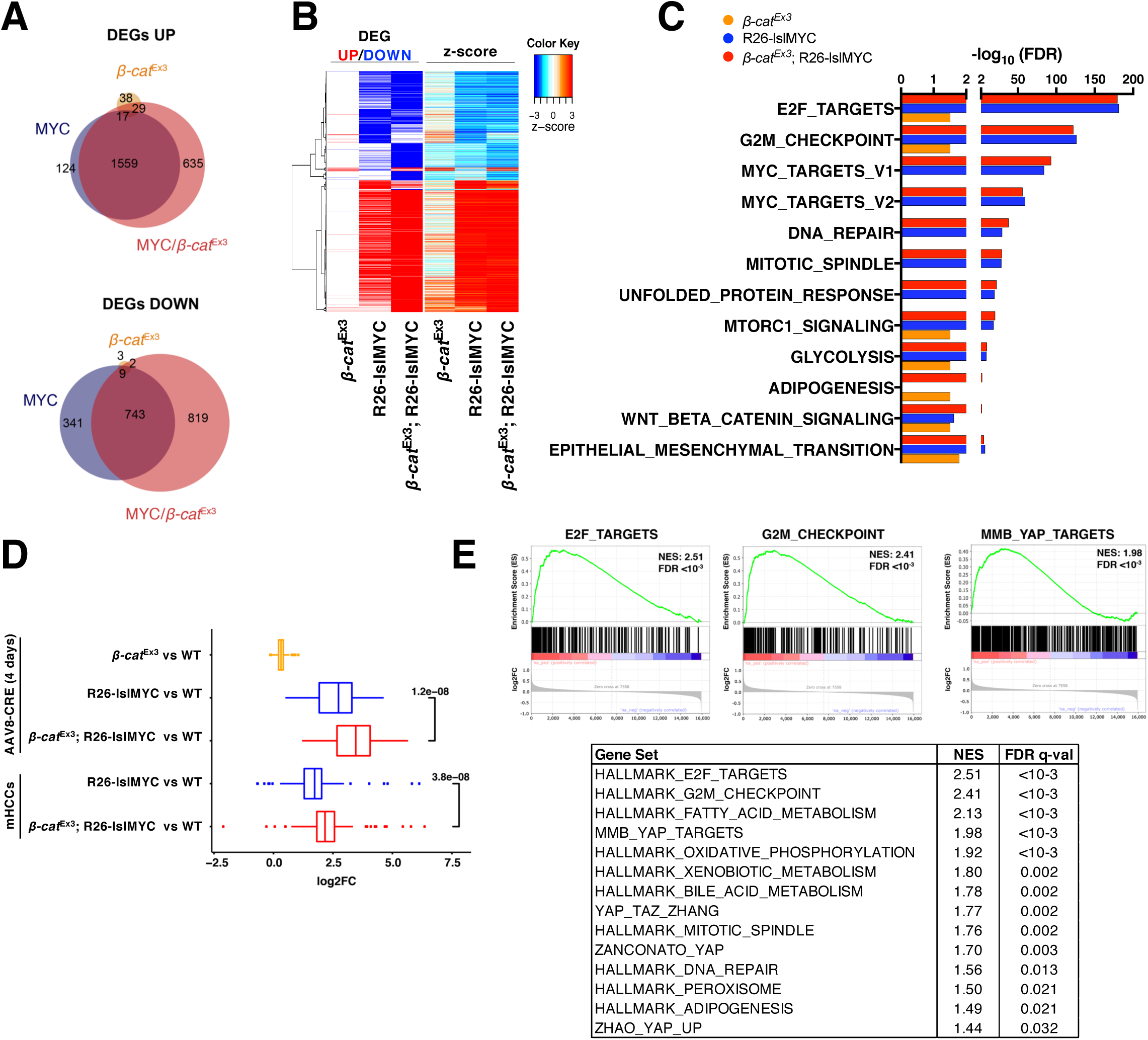
MYC and β-cat^Ex3^ promote the induction of a pro-proliferative Yap/Taz signature in the liver. RNA-seq profiling was performed in the livers of mice with activated R26-lslMYC and/or *β-cat*^Ex3^, and wild-type (wt) mice as controls. All samples were collected 4 days after high-titer AAV8-TBG-CRE injection. **(A)** Venn diagrams representing the overlap between differentially expressed genes (DEGs, qval <0.05) identified as induced (top) or repressed (bottom) relative to the wt control in each of the indicated groups. **(B)** Heatmap representation of the same DEGs. The left part (columns 1-3) shows up- and down-regulated genes, respectively marked in red and blue; the right part illustrates the z-score of the change in expression relative to wt for each DEG in each condition. **(C)** Bar-plots showing the False Discovery Rate (–log10) of the most significantly enriched gene ontology categories in at least one of the indicated genotypes. **(D)** Boxplot showing the fold-change in mRNA expression for the 125 genes of the MYC/β-catenin signature in each of the indicated conditions relative to the wt genotype (expressed as log2FC). p-values were calculated using Wilcoxon’s test. **(E)** Gene set enrichment plots for 3 examples (top) and complete list of Normalized Enrichment score (NES) and False Discovery Rate (FDR) values for all the significantly enriched hallmark gene sets (FDR<0.05) (bottom). Custom curated Yap/Taz signatures were added manually to the group of queried gene sets (see Methods for details). Genes were sorted from left to right according to the log2FC in their expression when comparing MYC/β-cat^Ex3^ versus MYC-overexpressing livers.

To address whether the cooperation between MYC and β-cat^Ex3^ in liver tumorigenesis may be associated with more subtle transcriptional changes, we focused on genes that were induced both by MYC and MYC/β-cat^Ex3^ overexpression, but whose fold change in expression relative to wild type livers was at least 1.5 times higher in MYC/β-cat^Ex3^ compared to MYC (Figure 2D): this led to the identification of a group of 125 genes that we will refer to as the “MYC/β-catenin signature”. Importantly, these genes were also induced in advanced mHCC tumors relative to wild type livers, and with a higher magnitude in MYC/β-cat^Ex3^ than in MYC-only tumors. Gene ontology analysis revealed that the MYC/β-catenin signature enriched - among others - for Yap/Taz transcriptional targets (Supplementary Table 3). Moreover, 59 out of 125 genes (*∼*45%) in our MYC/β-catenin signature (as opposed to *∼*14% of all active genes) scored as direct Yap targets by chromatin immunoprecipitation (28, 29) (Biagioni et al., manuscript in preparation) (Supplementary Table 3). Finally, Gene Set Enrichment Analysis (GSEA) revealed that several Yap/Taz signatures were also enriched in the transcriptional profiles of either MYC/β-cat^Ex3^ or MYC-overexpressing livers (Supplementary Figure 6B) and appeared, together with E2F targets, among the top enriched datasets when ranking all the genes by their fold-change between MYC/β-cat^Ex3^ and MYC-overexpressing livers (Figure 2E). Altogether, these data indicated that Yap and/or Taz are transcriptionally active in the liver upon MYC/β-cat^Ex3^ activation.

The enrichment of Yap/Taz-regulated genes in our dataset was particularly intriguing and, based on previous observations, might represent a possible mechanistic link between MYC and β-catenin: first, Yap/Taz are activated by WNT/β-catenin signaling owing to their association with the destruction complex (30); second, these factors cooperate with MYC in integrating mitogenic stimuli and promoting liver tumorigenesis (31); third, Yap/Taz promote proliferation by activating genes involved in either G1/S (28) or G2/M progression (29), consistent with the composition of our MYC/β-catenin signature. We thus focused on the possible involvement of Yap/Taz in the cooperative activity MYC and β-cat^Ex3^.

While the expression of Yap and Taz was barely detectable by immunohistochemistry analysis in normal liver, with the exception of cholangiocytes as previously reported (24), both proteins became detectable in a sizeable fraction of hepatocytes after AAV8-TBG-CRE-mediated activation of either MYC or β-cat^Ex3^ for 4-8 days, and were most strongly induced by concomitant activation of MYC and β-cat^Ex3^ (Figure 3A, B and Supplementary Figure 7A), paralleled by consistent changes in bulk protein levels (Figure 3C and Supplementary Figure 7B). In contrast, RT-qPCR analysis did not show any significant variation in *Yap/Taz* mRNA levels (Supplementary Figure 7C). In line with these observations, immunohistochemical analysis of established tumors confirmed increased expression of Yap and/or Taz relative to wild-type livers, albeit this occurred in either Myc-only or MYC/β-cat^Ex3^ mHCCs (Supplementary Figure 8). Finally, Taz (but not Yap) was induced and localized to the chromatin fraction also in in 3T9^MycER;S33Y^ fibroblasts upon induction of β-cat^S33Y^ (but in this system independently from MycER activation), correlating with increased expression of the canonical Yap/Taz transcriptional targets *Ctgf* and *Cyr61* (Supplementary Figure 9A and 9B).

**Figure 3.**
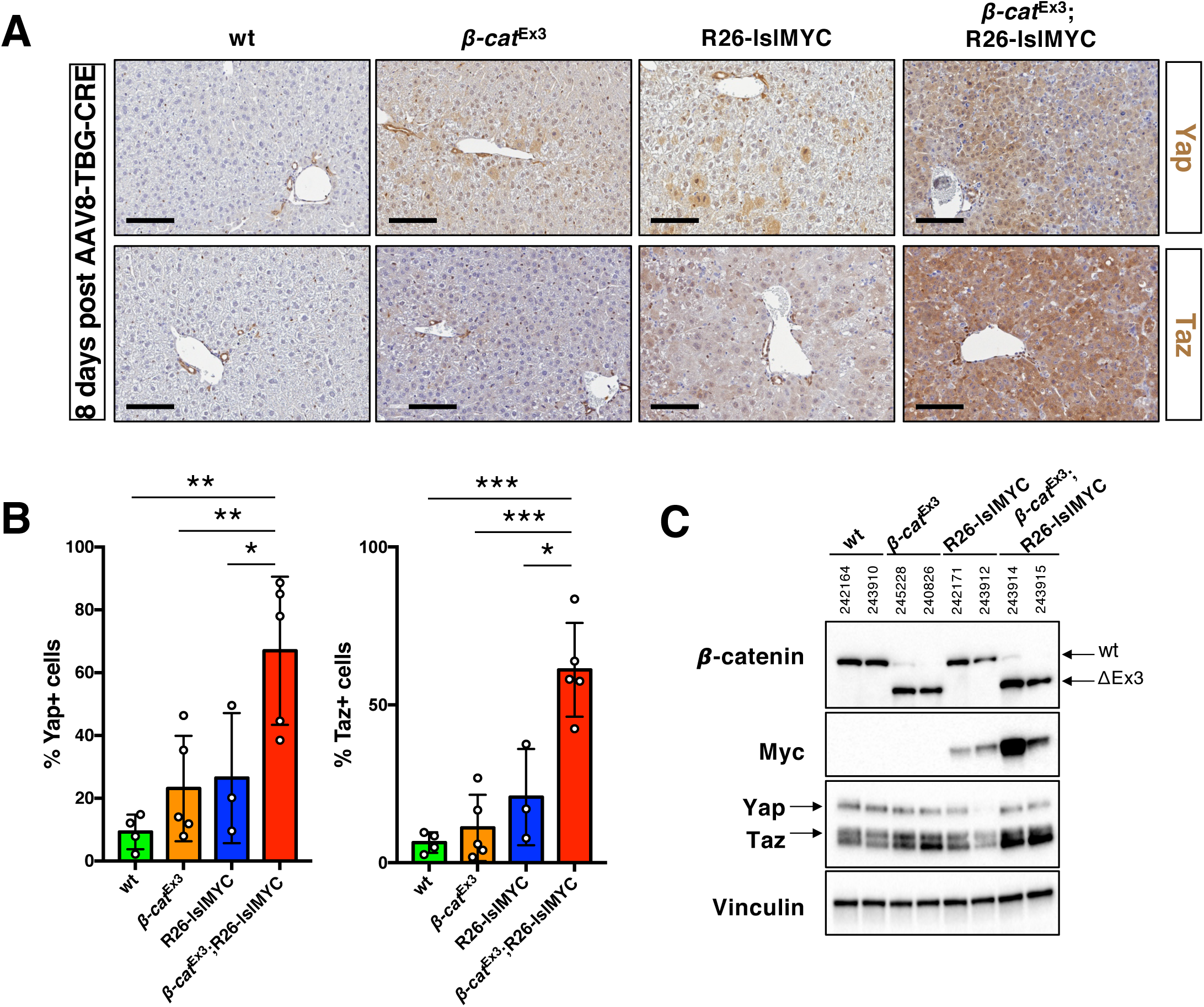
Increased Yap/Taz expression upon MYC and β-cat^Ex3^ activation in hepatocytes. **(A)** Immunohistochemical detection of Yap and Taz in representative liver samples from mice of the indicated genotypes, injected with high-titer AAV8-TBG-CRE 8 days prior to collection. Bar: 300 µm. **(B)** Quantification of the fraction of liver cells showing either cytoplasmic or nuclear positivity for Yap (left) and Taz (right) in mice of the different genotypes, 8 days after AAV8-TBG-CRE injection. Bar plot represent average and standard deviation for at least 3 biological replicates. **(C)** Representative Western blot analysis of Yap, Taz, β-catenin and MYC protein expression in lysates from livers sections from mice treated as in (**A**). Vinculin was used as loading control. Each sample is identified by its unique reference number.

Altogether our data support a model in which β-catenin promotes the stabilization of the Yap and Taz proteins, as previously reported (30), thus enhancing transcription of a subset of common Yap/Taz and MYC targets involved in promoting cell-cycle progression (31). In the liver, as opposed to fibroblasts, MYC also contributed to the up-regulation of Yap/Taz: while this effect remains to be explained at the mechanistic level, it further emphasizes the importance of Yap/Taz in Myc-dependent transformation (31).

### High expression of the MYC/β-catenin signature correlates with worst prognosis in HCC patients

Having defined a distinct MYC/β-catenin signature in mouse liver, we went on to address its significance in human cancer. In either of two independent HCC datasets (TCGA and LCI/FUDAN, see Supplementary Material & Methods), querying for the enrichment of our MYC/β-catenin signature allowed us to identify a distinct subgroup of patients whose tumors expressed high levels of the corresponding mRNAs (Figure 4A, B). In agreement with data from our mouse model, most of these patients also showed high expression of the MYC (32, 33), WNT (32, 33) and Yap/Taz (28) transcriptional signatures (Figure 4A, B, top; Fisher test on the TCGA cohort: odds ratio 192.90, p-value < 2.2e^-16^; on the LCI/FUDAN cohort: odds ratio 17.16346, p-value = 3.378e^-10^). Most relevant here, HCC patients with high level of expression of the MYC/β-catenin signature showed a significantly shorter survival compared with the ones with low expression (Figure 4C, D): in the TCGA dataset, where information about tumor grade was available (Figure 4A, top), the MYC/β-catenin signature also enriched for the most aggressive cases (grades 3 and 4, compared to grades 1 and 2; Fisher test: odds ratio 3.97, p-value = 5.5e^-08^).

**Figure 4.**
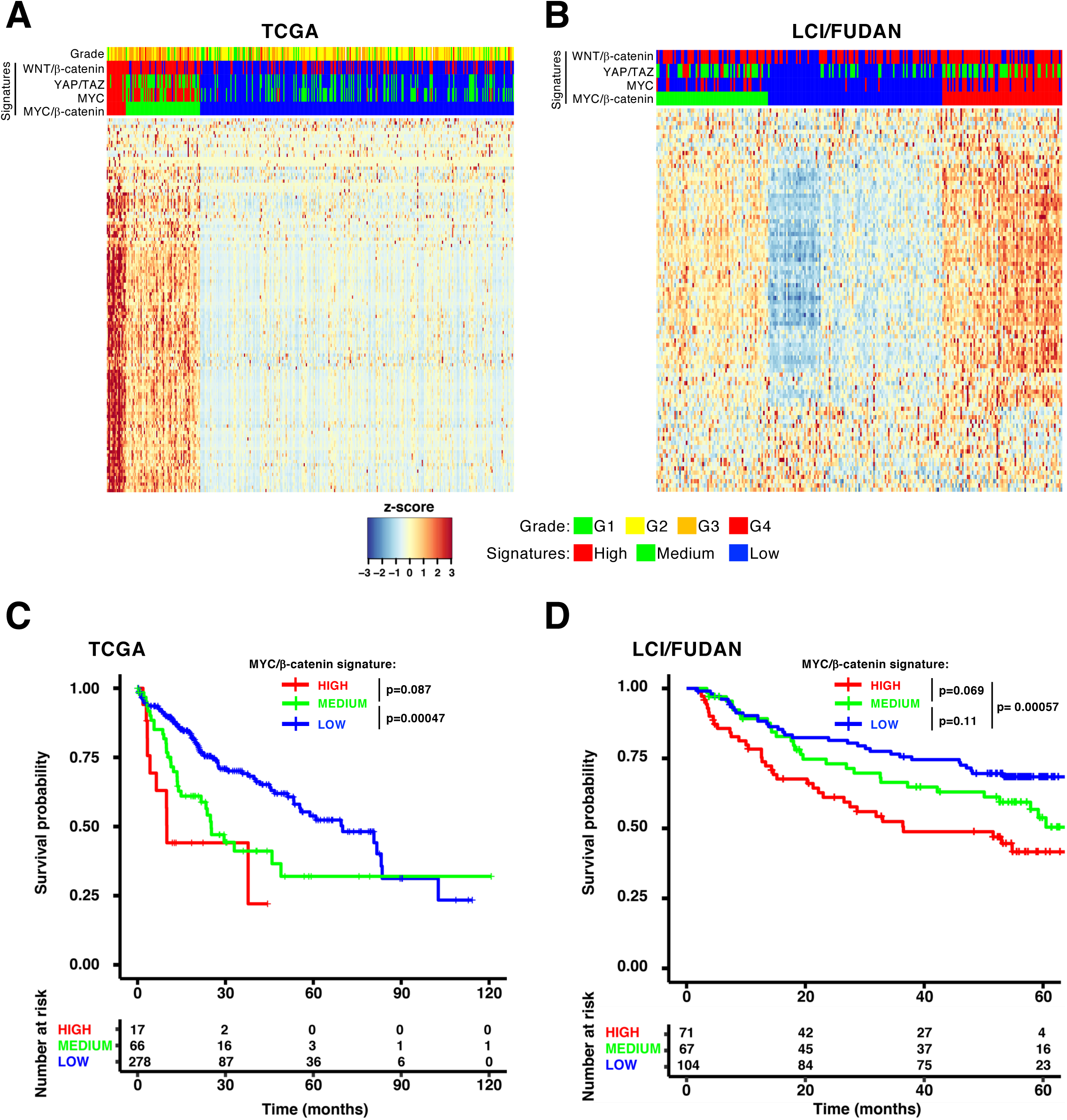
Clinical significance of the MYC/β-catenin signature in HCC. Patients in The Cancer Genome Atlas (TCGA, **A**) or LCI/FUDAN (**B**) cohorts were stratified according to the MYC/β-catenin gene expression signature. The level of expression of the MYC (HALLMARK_MYC_TARGETS_V1 from the Broad Institute repository), WNT/β-catenin (HALLMARK_WNT_BETA_CATENIN_SIGNALING from the Broad Institute repository) and YAP/TAZ (28) signatures for each sample is shown on top, with the addition of tumor grade for the TCGA cohort. The heatmap shows the z-score for the expression of each gene of the MYC/β-catenin signature in each patient. Kaplan-Meier plots showing the overall survival of HCC patients in the TCGA (**C**) and LCI/FUDAN (**D**) cohorts according to their level of expression of genes of the MYC/β-catenin signature. The indicated p-values were computed using the log-rank test. Number at risk: live patients in each cluster at the indicated time points.

### Yap/Taz are required for hepatocyte proliferation and tumor growth upon MYC/β-catenin activation

In order to address the role of Yap/Taz in MYC/β-cat^Ex3^-driven mHCCs, we generated a cohort of mice, in which Alb-CreER^T2^, β-cat^Ex3^ and R26-lsl-MYC were combined with conditional knockout alleles of Yap (34) and Taz (18) (hereafter Yap^f/f^;Taz^f/f^) and monitored those animals in the absence of TAM. Surprisingly, the presence of the Yap^f/f^;Taz^f/f^ alleles did not affect tumor development (Supplementary Figure 10A). However, PCR analysis showed that all tumors had retained at least one non-recombined Yap^f^ or Taz^f^ allele (Supplementary Figure 10B-C). Moreover, immunohistochemical analysis revealed expression of Yap and/or Taz in tumor sections (Supplementary Figure 10D). We surmise that Yap/Taz-null cells are counter-selected during tumorigenesis, pointing to an essential function of these proteins in MYC/β-cat^Ex3^-driven tumors. To corroborate this interpretation, we grew tumor cells *in vitro* and treated them with 4-OHT to activate CreER^T2^ and induce acute deletion of the remaining Yap^f^ or Taz^f^ alleles (Supplementary Figure 11A-E): as expected, this led to complete loss of the Yap and Taz proteins, while leaving MYC and β-cat^Ex3^ unaffected (Supplementary Figure 11D). In this setting, 4-OHT treatment caused a strong impairment in cell growth, which was not observed in cells bearing at least one wild-type allele of Yap and/or Taz (Figure 5A and Supplementary Figure 11E-G).

**Figure 5.**
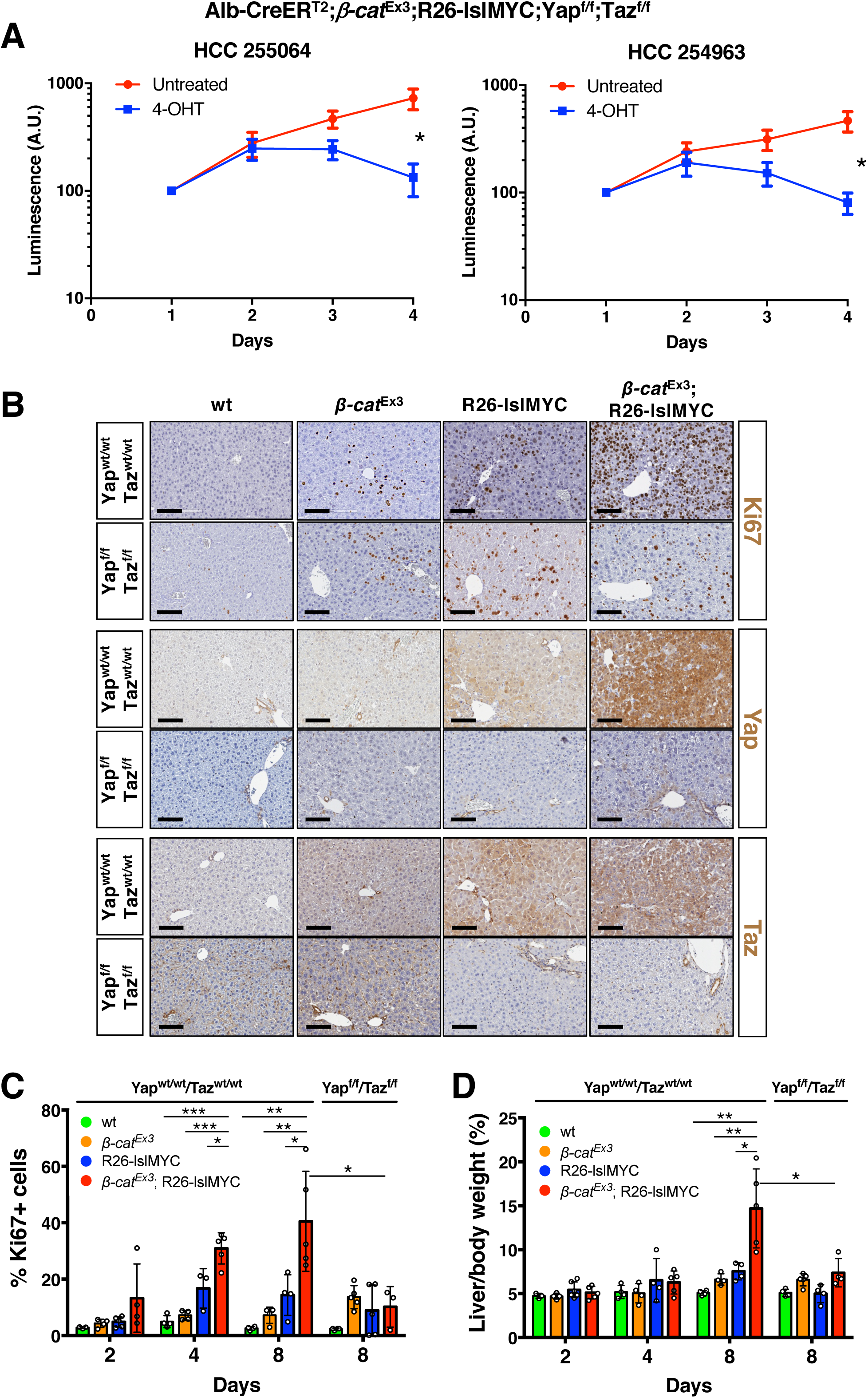
Yap/Taz are required for hepatocytes proliferation and liver tumorigenesis. **(A)** Growth curves for two independent Alb-CreER^T2^;*β-cat*^Ex3^;R26-lslMYC;Yap^f/f^;Taz^f/f^ mHCCs cultivated in the absence (red line) or presence (blue line) of 400nM 4-OHT to activate the recombinase activity of the CreER^T2^. Relative cell numbers were determined through Luminescence measurement with the Cell Titer Glo assay. A.U.: Arbitrary Units. **(B)** Immunohistochemical detection of Ki67 expression in representative liver sections from mice of the indicate genotypes, injected with AAV8-TBG-CRE 8 days prior to collection. Bar: 100 µm. **(C)** Fractions of Ki67-positive hepatocytes, and **(D)** Liver/body weight ratios (as %) for mice of the indicated genotypes, measured as in (B) at different time-points (2, 4 or 8 days) after AAV8-TBG-CRE injection. The bar plots represent average and standard deviation for at least 3 mice per group.

We then sought to address whether MYC and β-cat^Ex3^ might cooperatively induce proliferation when acutely induced in the liver, and whether this activity might be dependent upon Yap/Taz. To this aim, we combined the β-cat^Ex3^, R26-lslMYC, Yap^f/f^ and Taz^f/f^ alleles, infected mice with high-titer AAV8-TBG-CRE, and sacrificed animals at different time-point (2, 4, 8 days) to monitor proliferation (by Ki67 staining) and liver weight (Figure 5B-D): mice co-expressing MYC and β-cat^Ex3^ (with wild-type Yap and Taz) showed increased hepatocyte proliferation over the time-course, preceding a marked increase in liver/body weight ratio at day 8. Strikingly, these responses were abrogated by deletion of Yap/Taz (Figures 5B-D). It is noteworthy here that WNT-1 overexpression and β-catenin activation were reported to dampen MYC-induced apoptosis in rat fibroblasts and intestinal epithelial cells (35). However, this mechanism was not verified in our model: as judged by TUNEL staining, MYC-induced apoptosis was not decreased, but rather increased in the presence of β-cat^Ex3^ (Supplementary Figures 12A, B), most likely associated with the hyper-proliferative response.

Altogether our results support an essential, but mutually redundant role for Yap/Taz as downstream effectors of β-cat^Ex3^, driving the cooperation with MYC through activation of a set of genes, which in turn promote proliferation in hepatocytes.

## Discussion

In this study, we describe a mouse model that recapitulates the co-activation of MYC and WNT/β-catenin observed in a fraction of human HCCs (2, 3). While confirming that Myc is not a downstream transcriptional target of WNT/β-catenin in the liver (9), as opposed to other tissues (5–8), our data revealed that concomitant activation of both pathways in mouse hepatocytes promoted unscheduled cell proliferation and cooperated in the onset of HCC-like tumors. This cooperativity between MYC and WNT/β-catenin can be extended to other tissues, as reported in T-cells (12) and as verified here in immortalized fibroblasts. While our work was under completion, others reported a cooperative action of Myc and β-catenin in liver tumorigenesis, based on hydrodynamic-tail vein injection of plasmids expressing both oncogenes (21). These authors showed that the Myc/β-catenin cooperation relies on the ability of β-catenin to promote immune escape: in particular, β-catenin suppressed expression of the chemokine Ccl5, resulting in defective recruitment of dendritic cells and impaired T-cell activity (21). Consistent with this scenario, our MYC/β-cat^Ex3^ mHCCs showed a slight reduction in *Ccl5* mRNA levels compared to MYC-driven mHCCs, while short term activation of MYC and/or β-catenin had no significant impact on *Ccl5* (data not shown). While the possible contribution of immune-surveillance - or other systemic effects - remains to be investigated in our model, our results demonstrate that MYC and β-catenin have a direct, cell-autonomous effect on cellular transformation.

In order to address the mechanisms underlying the cooperativity of MYC and β-catenin, we profiled gene expression shortly after their co-activation in hepatocytes. Our data did not reveal any major interference between MYC- and /β-catenin-regulated transcriptional programs, but unveiled the existence of a discrete MYC/β-catenin signature of 125 genes, which were more induced upon co-activation of both oncogenes and mainly encoded products involved in cell cycle control and proliferation. Strikingly, this signature enriched for genes previously reported to be under the direct control of the transcriptional co-activators Yap/Taz. In subsequent experiments, we showed that Yap and Taz were induced upon short-term activation of MYC and β-cat^Ex3^ in the liver, and were essential for the proliferative response of hepatocytes. In line with these observations, Yap/Taz were also abundant in MYC/β-cat^Ex3^ tumors and were required for the growth and survival of tumor cells.

Our findings on the role of Yap/Taz as effectors in the cooperative action of MYC and β-catenin connects two apparently unlinked regulatory cross-talks, including (*i.)* the activation of Yap/Taz by WNT/β-catenin signaling (18, 30, 36) and (*ii.)* the cooperation between Myc and Yap in supporting either cell proliferation in serum-stimulated fibroblasts, or tumorigenesis in the liver (31). First, besides signaling cues such as the Hippo pathway, mechanotransduction (37) or a non-canonical (β-catenin-independent) WNT pathway (38), Yap/Taz have been linked to WNT/β-catenin signaling (18, 30, 36). Taz in particular was shown to be recruited to - and degraded by the destruction complex in a β-catenin-dependent manner (30). Moreover, Yap/Taz were stabilized and activated following either knock-out of Apc in mouse tissues and human cell lines (18, 30, 36), or expression of the stable β-catenin mutant β-cat^Ex3^ in the small intestine (36). It is noteworthy here that additional context-dependent mechanisms may be involved in the aforementioned effects: in fibroblasts, activation of β-cat^S33Y^ (but not Myc) induced accumulation of Taz; in hepatocytes, instead, either β-cat^Ex3^ or Myc alone caused mild increases in Yap/Taz levels, which were markedly enhanced by the co-activation of both oncogenes. Altogether, while the molecular basis for these additional effects remains to be addressed, the above findings establish Yap/Taz as downstream effectors of WNT/β-catenin signaling. As a second connection, Yap was shown to cooperate with Myc in supporting either cell proliferation or tumorigenesis, and did so through the joint activation of a subset of proliferation-associated genes (31). Indeed, almost half of the genes included in our MYC/β-catenin signature were independently identified as direct targets of Yap, and were induced upon co-activation of MYC and Yap in the liver. Altogether, we conclude that Yap/Taz link WNT/β-catenin activation to a MYC-regulated proliferative program, underlying the cooperativity between these two oncogenic pathways.

While the involvement of MYC, WNT/β-catenin and Yap/Taz in liver tumorigenesis was amply documented (3, 13, 31, 39–43), their mutual interplay – if any – remained to be unraveled. Toward this aim, we re-analyzed gene expression profiles from two independent patient cohorts, and observed that the MYC/β-catenin signature identified in our work was enriched in a subset of HCC patients, in which it correlated with worse prognosis.

In recent years, Yap/Taz have emerged as key player in tumorigenesis in different tissues (37), and several approaches for the inhibition of their expression and/or activity have been proposed as potential therapeutic options (44). Our results add a new element in this regard, indicating that a therapeutic strategy aimed at tackling Yap/Taz may be particularly effective in tumors with aberrant activation of MYC and WNT/β-catenin. Moreover, targeting Yap and Taz may be therapeutically safe, given that their deletion showed limited short-term effects in normal tissues, including the liver (34, 45). In this regard, an attractive class of molecules – already in use in the clinic – are statins, which have been demonstrated to (i) suppress Yap/Taz function by impairing their nuclear import and promoting their degradation (46, 47), (ii) reducing Taz-dependent HCC proliferation (48) and (iii) decreasing the risk of cancer mortality, in particular for HCC (49, 50).

Altogether, our work in mice has revealed a novel functional interplay between MYC, WNT/β-catenin and Yap/Taz in liver tumorigenesis. Most importantly, these interactions hold true in human HCC, with important prognostic implications. Future work will address whether targeting these pathways may hold personalized therapeutic potential in patients whose tumors show co-activation of these pathways.

## Disclosure of Potential Conflicts of Interest

The authors declare that they have no conflict of interest.

## Authors’ Contributions

AB, GPGF, GB, FB, GC, NT, FC and AS performed experiments. MD provided technical support. DO conducted pathological analyses. MF, MJM, VP and AS performed bioinformatic data analysis. DP supervised the work of FC. AB, SC, AS and BA designed experiments, supervised the work, and wrote the manuscript.

## Acknowledgements

Acknowledgments

We thank Gioacchino Natoli, Ottavio Croci, Thomas Valenta, Konrad Basler and members of the Amati’s and Campaner’s labs for discussion and advice. We also thank Konrad Basler, Eduard Battle, Pierre Chambon, Stefano Piccolo, Makoto Taketo, Thomas Valenta for materials, Camilla Recordati for initial pathological analyses, A. Gobbi, M. Capillo and all the members of the animal facility for their help with the management of mouse colonies, S. Bianchi, L. Rotta, and T. Capra for assistance with Illumina sequencing.

## Supplementary Material and Methods

### Genotyping of mice

Genomic DNA was extracted from tail biopsies by overnight digestion in lysis buffer (100 mM TrisHCl pH 8. 5,5 mM EDTA pH 8, 0, 2% SDS, 0,2 M NaCl and 0,1 mg/ml freshly added Proteinase K) at 55°C, followed by heat inactivation (5 minutes at 100°C) and dilution with 10 volumes of water, and then used to evaluate offspring genotype by semi-quantitative PCR (GoTaq® G2 Hot Start Polymerase, #M7408). The primers used for each genotyping are listed in Supplementary table 4.

### Cell lines

3T9^MycER^ fibroblasts^1^ and 293T cells were grown in DMEM medium supplemented with 10% serum, 1% penicillin/streptomycin and 2 mM L-Gln. To generate the 3T9^MycER;S33Y^ cells, the cells were infected with a doxycycline-inducible pSLIK lentiviral vector (https://www.addgene.org/25737/) encoding mouse β-catenin S33Y (a kind gift of T. Valenta and K. Basler)^2^, selected with Hygromycin B (ThermoFisher Scientific, #10687010) and kept in culture in 3T9^MycER;S33Y^ medium (10% tet-free FBS - Fetal bovine serum EU TET free Euroclone, #ECS0182L, 1% penicillin/streptomycin and 2 mM L-Gln).

### Culture of primary mHCC tumor cells

mHCC cells were isolated and cultured accordingly to published protocols^3^, with the following modifications. Briefly, individual tumor nodules were minced to small pieces with a scalpel and digested for 30 minutes at 37°C with gentle shaking in 50 ml/tumor of DMEM medium, freshly supplemented with 0.5 U/ml dispase (Stemcell technologies #07913) and 0.1 U/ml collagenase (Sigma-Aldrich, #C2674). Cells were passed 3 times through a 70 μm nylon mesh filter (Falcon #352350), centrifuged at 80g for 10 minutes and resuspended in 10 ml of Erythrocyte Lysis buffer (150 mM NH_4_Cl, 10 mM KHCO_3_, 0.1 mM EDTA). After a centrifugation step at 335g for 5 minutes, the remaining cells were resuspended in 30 ml of medium and plated in a 10 cm dish for 1 hour at 37°C in medium to eliminate fibroblasts attaching to the plastic. The cells present in the supernatant were centrifuged at 335g for 5 minutes and then plated in mHCC medium (DMEM, 10% FBS, 1% penicillin/streptomycin and 2 mM L-Gln, 40 ng/mL murine HGF (Peptrotech, #315-23), 1 μM Dexametasone (Sigma-Aldrich, #D4902) and 20 ng/mL EGF (Peprotech, #100-15) on culture dishes previously coated with 0.1% Gelatin (Sigma-Aldrich, #G2500).

### Western Blot

Liver tissue was homogenized with gentleMACS M Tubes (Miltenyi Biotech, 130-096-335) and the gentleMACS™ Dissociator (Miltenyi Biotech, #130-093-235) in Western Blot lysis Buffer (300 mM NaCl, 1% NP-40, 5 mM Tris pH 8, 0.1 mM EDTA) freshly supplemented with protease (cOmplete™, Mini Protease Inhibitor Cocktail, Roche-Merck, #11836153001) and phosphatase inhibitors (PhosSTOP, Roche-Merck, #4906845001). In vitro cultured mHCCs and 3T9^MycER;S33Y^ were washed twice in PBS and then scraped in Western Blot lysis Buffer. Cell or liver lysates where then sonicated for 10 seconds in a Branson Sonifier 250 (Output Control = 2) equipped with a 3.2 mm Tip (Branson, #101-148-063), cleared by centrifugation at 16000g for 15 minutes at 4°C and quantified by Bradford assay (Bio-Rad Protein Assay, #5000006). After addition of 6X Laemmli buffer (375 mM Tris-HCl, 9% SDS, 50% glycerol, 9% beta-mercatoethanol and 0.03% bromophenol blue), lysates were boiled (5 minute at 100°C) and then electrophoresed on pre-cast 4-15% gradient Polyacrilamide gels (Bio-Rad, #5678084), transferred onto methylcellulose membranes (Bio-Rad, #1704271) and protein expression detected with the indicated primary antibodies (Supplementary Table 5). Chemiluminescence was detected using a CCD camera (ChemiDoc XRS+ System, Bio-Rad). Quantification of protein levels was performed using the Image Lab software (Bio-Rad, version 4.0).

### Colony formation assay

Colony formation was assessed as follows: 500 3T9^MycER;S33Y^ fibroblasts were plated in triplicate in non-treated 24-well plates (Costar, #3738) in 2 ml/well of a solution composed of 50% (v/v) methylcellulose (MethoCult™ SF, Stemcell Technologies, #M3236) and 50% (v/v) medium. Where applicable, 400 nM of 4-hydroxytamoxifen (OHT; Sigma-Aldrich #H7904) and/or 1 μM doxycyline hydrate (Sigma-Aldrich, #D9891-100G) were added. Colonies of at least 10 μM of diameter were counted 7-10 days after plating.

### Subcellular fractionation

For subcellular fractionation, ∼5*10^6^ 3T9^MycER;S33Y^ fibroblasts were trypsinized and pelleted (335g, 5 min, 4°C). After one wash with PBS, cells were resuspended in 200 μl of pellet of RSBS buffer (10 mM Tris/HCl pH 7.5, 10 mM NaCl, 5 mM MgAcetate) and incubated for 10 minutes at 4°C. NP-40 (CAS 9016-45-9; Merck, #492016-100ML) was added to a final concentration of 0.2% and the integrity of nuclei checked by trypan blue staining. At this stage 20 μl of each sample (corresponding to 1/10 of the total volume) were collected and supplemented with 6X Laemmli buffer. This represents the whole-cell lysate. The remaining material was centrifuged at 335g for 10 minutes at 4°C to pellet nuclei, and the supernatant (∼150 μl) collected and supplemented with 6X Laemmli buffer (Cytosolic fraction). Nuclei were then washed twice by gentle resuspension with 1 ml of Washing Buffer (0.88 mM Sucrose, 5 mM MgAcetate), centrifuged for 5 minutes at 380g at 4°C, and lysed by resuspension in 150 μl (a volume equal to the supernatant collected for the cytosolic fraction) of a solution composed of one half of Glycerol Buffer (20 mM Tris/HCl pH 8, 75 mM NaCl, 0.5 mM EDTA, 50% glycerol, 0.85 mM DTT) and one half of Nuclei Lysis Buffer (20 mM Hepes pH 7.6, 7.5 mM MgCl2, 0.2 mM EDTA, 300 mM NaCl, 1 M UREA, 1% NP40, 0.1 mM DTT), brief vortexing and incubation at 4°C for 10 minutes. This nuclear lysate was then centrifuged at 16000g for 2 minutes at 4°C, the supernatant collected (∼150 <u>μ</u>l) and supplemented with 6X Laemmli buffer (Nucleoplasmic fraction). The remaining pellet (Chromatin-associated fraction) was washed twice in Glycerol Buffer/Nuclei Lysis Buffer, resuspended in ∼180 μl of 1X Laemmli Buffer and sonicated for 10 seconds in a Branson Sonifier 250 (Output Control = 2) equipped with a 3.2 mm Tip (Branson, #101-148-063). All buffers were freshly supplemented with protease (cOmplete™, Mini Protease Inhibitor Cocktail, Roche-Merck, #11836153001) and phosphatase inhibitors (PhosSTOP, Roche-Merck, #4906845001) before usage. Equal volumes of each fraction were then analysed by Western Blotting with the appropriate antibodies (Supplementary Table 5).

### Immunohistochemistry

Freshly isolated liver tissue was washed in PBS, fixed in 4% (v/v) paraformaldehyde at 4°C degrees for at least 16-24 hours, washed in PBS, and then stored in 70% ethanol at 4°C until further processing. The tissue was dehydrated with increasing concentrations of ethanol, embedded in paraffin blocks, cut into 3/5-mm thick sections and mounted on glass slides. Sections were dewaxed and rehydrated through an ethanol scale, heated in citrate solution (BioGenex, #HK086-9K) in a water bath at 99°C for 30 minutes for antigen unmasking, washed once in water, and treated with 3% H_2_O_2_ for quenching of endogenous peroxidases. After overnight incubation at 4°C with the relevant primary antibody (Supplementary table 5), slides were washed twice with TBS and incubated with secondary antibodies (Supplementary Table 5) for 45 minutes. The signal was revealed with DAB peroxidase substrate solution (Dako, #K3468) for 1 to 5 minutes. Slides were finally counterstained with Harris Hematoxylin (Sigma-Aldrich, #HHS80), dehydrated through alcoholic scale, and mounted with Eukitt (Bio-Optica, #09-00250).

Apoptosis was measured on FFPE liver sections derived from mice of the different genotypes with a TUNEL Assay Kit (Abcam, ab206386) according to the manufacturer’s instructions, with the exception of counterstaining, which was performed with Harris Hematoxylin.

All images were acquired with the Aperio Digital Pathology Slide Scanner ScanScopeXT (Leica) and the percentage of positive cells and signal intensity were quantified with the Aperio ImageScope software (Leica).

### RNA extraction and RT-qPCR analysis

Liver tissue was homogenized with GentleMACS M Tubes (Miltenyi Biotech, #130-096-335) and the gentleMACS™ Dissociator (Miltenyi Biotech, #130-093-235) in RNA lysis buffer (Zymo, #R1054), while *in vitro* cultured mHCC and 3T9^MycER;S33Y^ cells were scraped in PBS, pelleted and then resuspended in RNA lysis buffer. Total RNA was purified from these lysates onto Quick-RNA Miniprep columns (Zymo, #R1054) and treated on-column with DNaseI. Complementary DNA (cDNA) was prepared using the ImProm-II^TM^ reverse transcription kit (Promega, #A3800). 10 ng of cDNA were used as template for each real-time PCR reaction. cDNA was detected by fast SyberGreen Master Mix (Applied Biosystems, #4385614) on a CFX96 Touch™ Real-Time PCR Detection System (Biorad). The PCR primers used are listed in Supplementary Table 4.

For RNA-seq experiments, total RNA was purified as above, and RNA quality was checked with the Agilent 2100 Bioanalyser (Agilent Technologies). 0.5-1 μg were used to prepare libraries for RNA-seq with the TruSeq stranded total RNA Sample Prep Kit (Illumina, #20020596) following the manufacturer’s instructions. RNA-seq libraries were then run on the Agilent 2100 Bioanalyser (Agilent Technologies) for quantification and quality control and pair-end sequenced on the Illumina 2000 or NovaSeq platforms.

### Next generation sequencing data filtering and quality assessment

RNA-seq reads were filtered using the fastq_quality_trimmer and fastq_masker tools of the FASTX-Toolkit suite (http://hannonlab.cshl.edu/fastx_toolkit/). Their quality was evaluated and confirmed using the FastQC application (https://www.bioinformatics.babraham.ac.uk/projects/fastqc/). Pipelines for primary analysis (filtering and alignment to the reference genome of the raw reads) and secondary analysis (expression quantification, differential gene expression) have been integrated in the HTS-flow system^4^. Bioinformatic and statistical analyses were performed using R with Bioconductor and comEpiTools packages^5, 6^.

### RNA-seq data analysis

RNA-seq NGS reads were aligned to the mm9 mouse reference genome using the TopHat aligner (version 2.0.8)^7^ with default parameters. In case of duplicated reads, only one read was kept. Read counts were associated to each gene (based on UCSC-derived mm9 GTF gene annotations), using the featureCounts software (http://bioinf.wehi.edu.au/featureCounts/)^8^ setting the options -T 2 -p -P. Differentially expressed genes (DEGs) were identified using the Bioconductor Deseq2 package^9^ as genes whose q-value is lower than 0.05.

The MYC/β-catenin signature has been identified using the following criteria:

qval (R26-lslMYC vs wt) <0.05
qval (*β-cat*^Ex3^;R26-lslMYC vs wt) <0.05
log2FC (R26-lslMYC vs wt) >0
log2FC (*β-cat*^Ex3^;R26-lslMYC vs wt) - log2FC (R26-lslMYC vs wt) >0. 5

Functional annotation analysis to determine enriched Gene Ontology categories was performed using the online tool at http://software.broadinstitute.org/gsea/index.jsp. Gene set enrichment analysis (GSEA) was performed using the Desktop tool of the Broad Institute (http://software.broadinstitute.org/gsea/index.jsp) with the addition of the following manually curated lists of Yap/Taz targets:

- Yap_Taz_ZHANG: genes commonly induced by Yap and Taz from the supplementary Table 1 of reference^10^
- ZHAO_Yap_UP: Yap upregulated genes, from the supplementary Table 1 of reference^11^
- CORDENONSI_Yap_CONSERVED_SIGNATURE^12^: from the Broad Institute repository (tool http://software.broadinstitute.org/gsea/index.jsp)
- ZANCONATO_Yap: Yap/Taz/TEAD direct targets, from supplementary Table 3 of reference^13^
- MMB_Yap_TARGETS, Yap_DIRECT_0. 5KB, as defined in^14^; lists obtained from the authors
- Dupont_Yap: Yap upregulated genes, from the supplementary Table 1 of reference^15^

### Statistical analysis

All the experiments, were performed at least in biological triplicates. Sample size was not predetermined, but is reported in the respective Figure legends. Two-tailed Student’s t-test was used to compare between two groups and expressed as p-values. In all Figures: *p < 0.05, **p < 0.01, ***p < 0.001, ****p<0.0001

### Survival analysis

RNA-seq samples and patient’s survival data for LIHC (HCC) were downloaded from the TCGA Data Portal (https://tcga-data.nci.nih.gov/tcga/) and used for survival analyses. TCGA RNA FastQ files were processed with the same pipeline used for RNA-seq data except for being aligned to hg19. The data were first normalized to Transcripts Per kilobase Million (TPM) then, for each gene, the z-score across all samples was calculated. The affinity propagation clustering (apcluster R package)^16, 17^ was used to group samples by expression of the genes composing the signatures. For each cluster, the median value of z-scores was computed. Then clusters were ordered accordingly and labeled as “high”, “medium” or “low”. The Log-rank test was applied (survminer R package, https://rpkgs.datanovia.com/survminer/index.html) for comparing survival curves from patients belonging to different clusters (https://CRAN.R-project.org/package=survival)^18^. Only samples from primary tumors were considered.

An independent cohort of 242 HCC patients from Liver Cancer Institute and Zhongshan Hospital, Fudan University^19^ was used for validation. In this case, patient’s tumor samples were profiled for gene expression with Affymetrix GeneChip HG-U133A 2.0 arrays. Probes were matched to the correspondent gene and for genes with more than 1 probe set, the mean gene expression was calculated. The downstream analysis applied is the same as for the TCGA dataset.

### Oligonucleotide Primers

Primers for mRNA analysis were designed with Primer-BLAST (https://www.ncbi.nlm.nih.gov/tools/primer-blast/)^20^. The complete list of primers used in this study is shown in Supplementary Table 4.

### Code availability

All R scripts used in data analysis and generation of Figures are available upon request.

### Data availability

RNA-seq data have been deposited in NCBI’s Gene Expression Omnibus (GEO) and are accessible through GEO series accession number GSE138296.

---

**Supplementary Figure 1.**
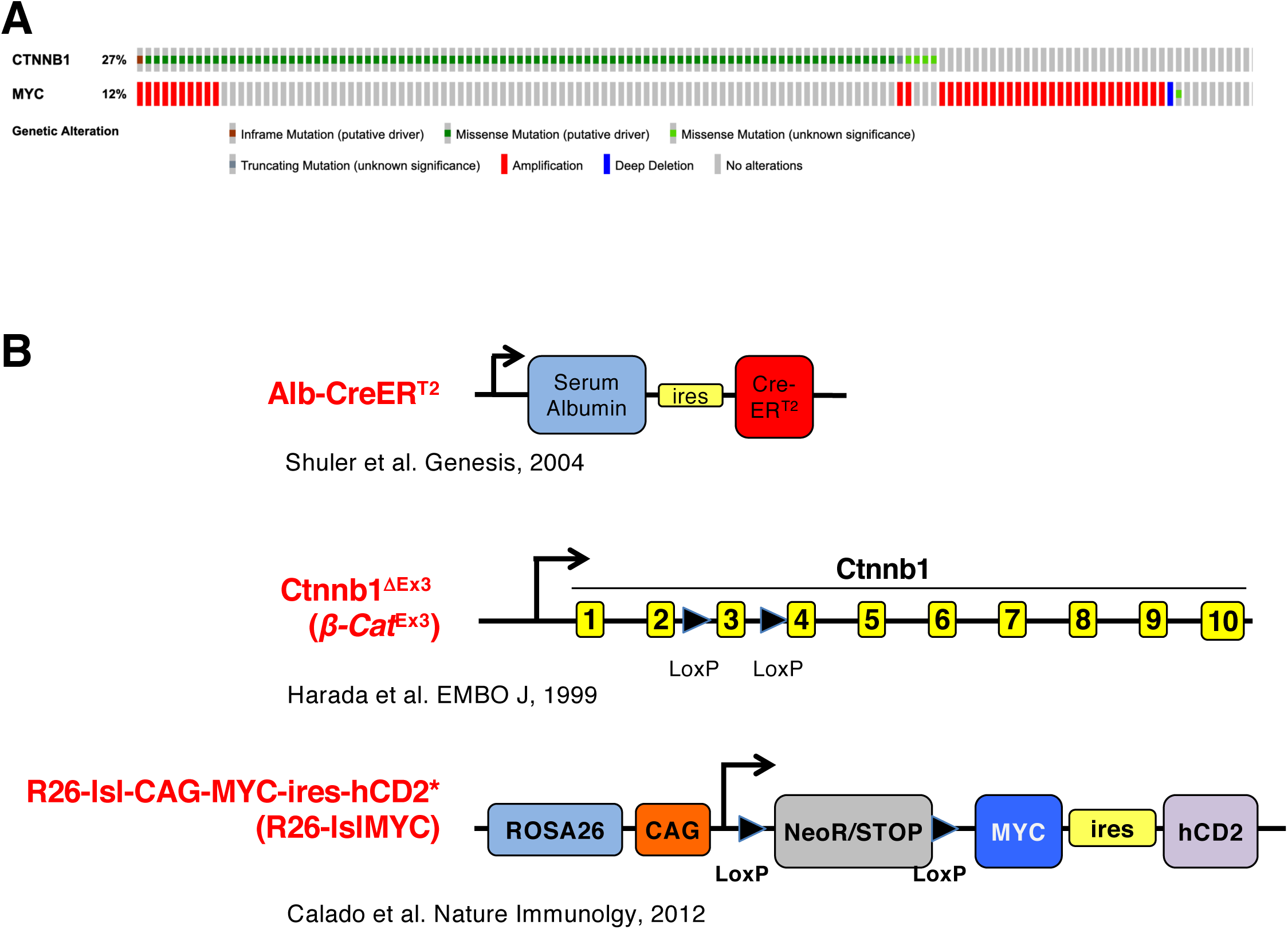
A new mouse model of liver tumors modeling MYC and β-cat^Ex3^ deregulation observed in human HCCs. **(A)** Oncoprint of *MYC* and *CTNNB1* alterations in 366 HCC patients (TCGA, provisional, December 2018, cbioportal). **(B)** Schematic overview of the recombinant mouse alleles used in this study.

**Supplementary Figure 2.**
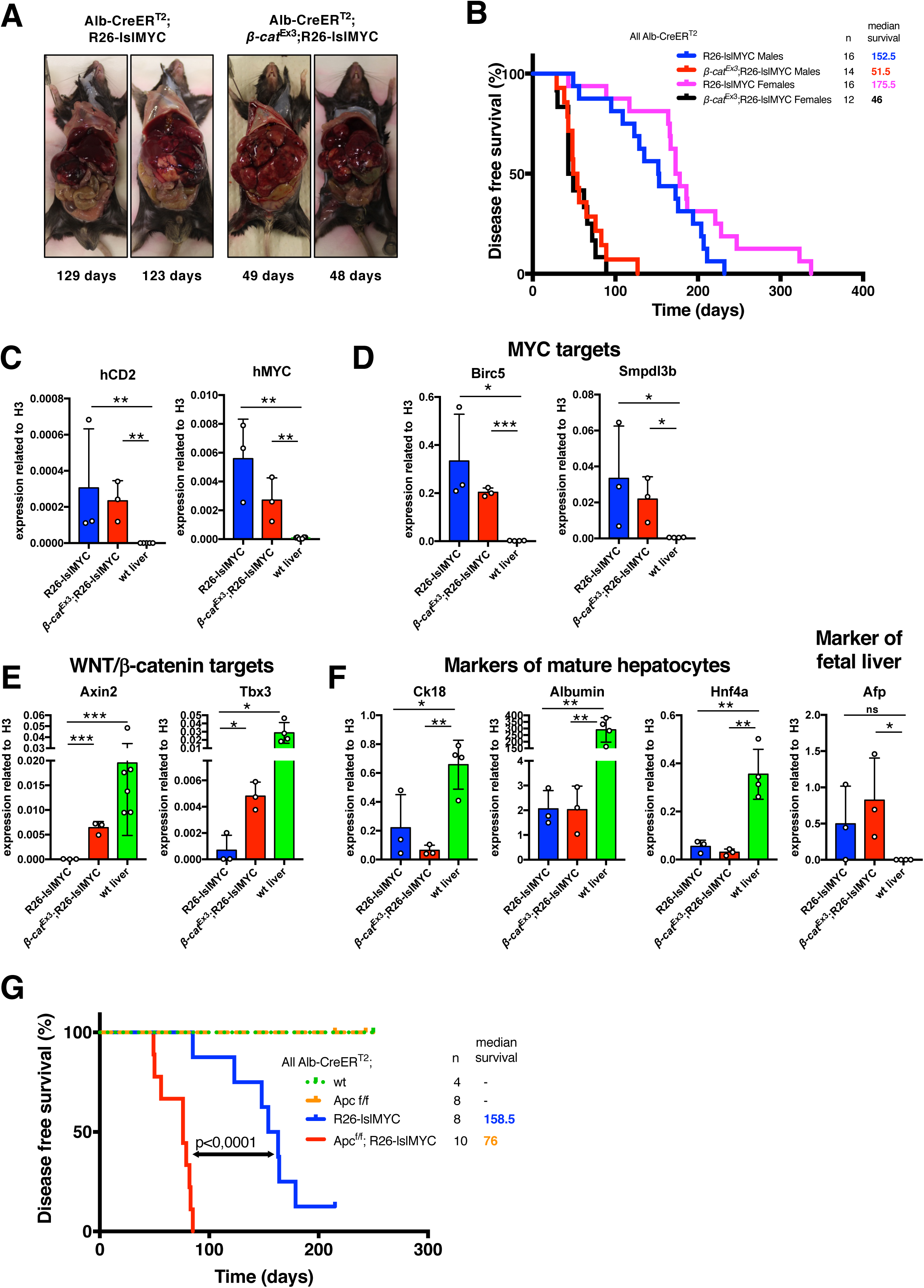
MYC and β-cat^Ex3^ cooperate in liver tumorigenesis. **(A)** Representative pictures of liver tumors in Alb-CreER^T2^;R26-lslMYC and Alb-CreER^T2^;*β-cat*^Ex3^;R26-lslMYC mice, kept Tamoxifen-free over their lifetime. The age (days) at which each animal was sacrificed is indicated. **(B)** Kaplan Meyer disease-free survival curves for mice of the indicated genotypes: the animals are the same as in Figure 1A, but are divided here between males (M) and females (F). The number of mice (n) and the median survival are indicated. p-values were calculated with the log-rank test. **(C-F)** RT-qPCR analysis of selected mRNAs in tumors of the indicated genotypes (all in the presence of the Alb-CreER^T2^ transgene), with wild-type liver as control: **(C)** human MYC and CD2, encoded by the R26-lslMYC allele (see Supplementary Figure 1A); **(D)** Representative MYC-activated genes; **(E)** Representative WNT/β-catenin-activated genes; **(F)** Representative markers of mature or fetal hepatocytes. (F) in tumors of the indicated genotypes or wild-type liver as control. Average values and standard deviations were derived from at least 3 biological replicates. **(G)** Deletion of *Apc* also accelerates MYC-driven carcinogenesis: Kaplan Meyer disease-free survival curves for mice of the indicated genotypes (all in the presence of the Alb-CreER^T2^ transgene). The number of mice (n) and the median survival are indicated. p-values were calculated with the log-rank test.

**Supplementary Figure 3.**
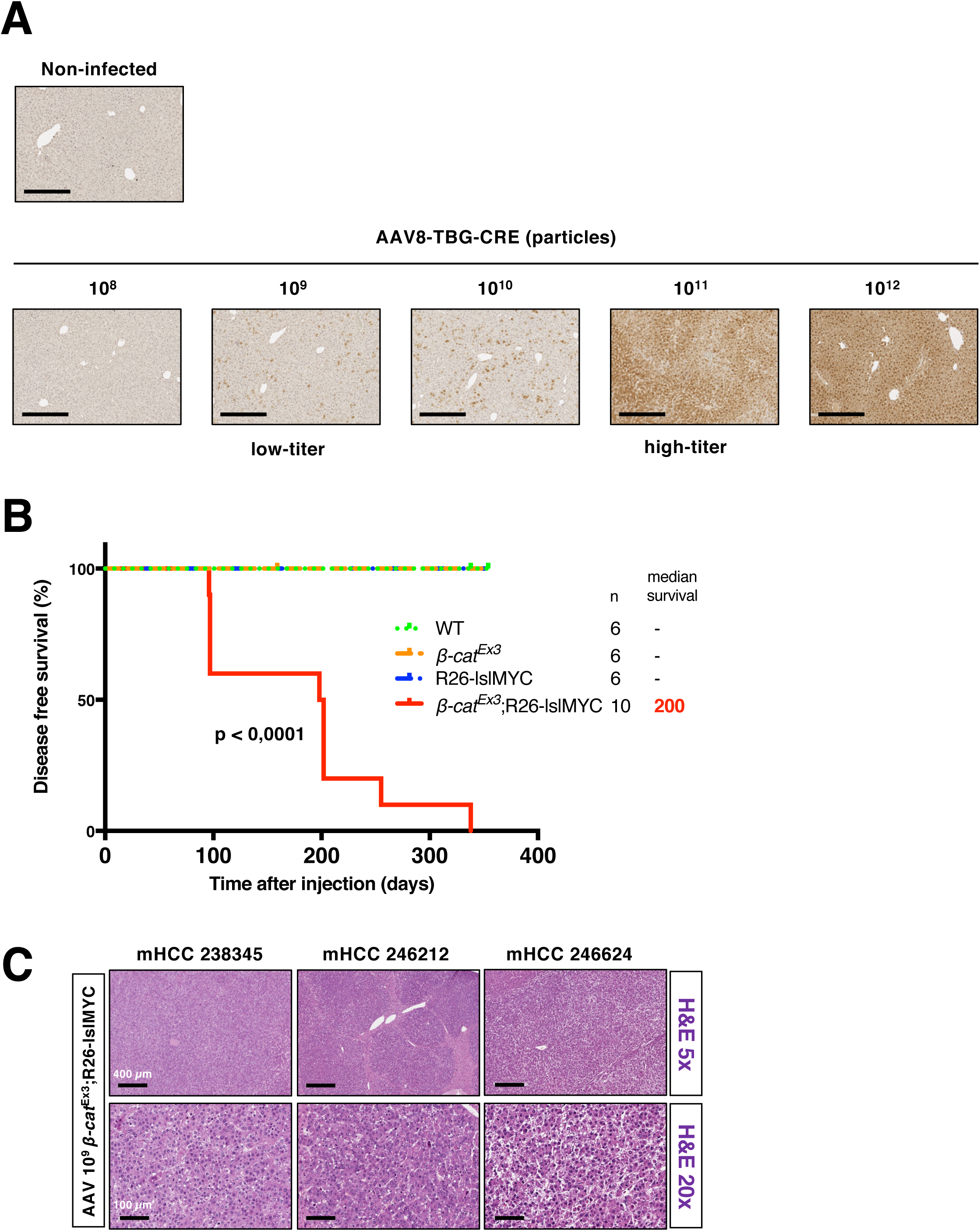
β-cat^Ex3^ activation by AAV8-TBG-CRE also accelerates MYC-driven tumorigenesis. **(A)** Immunohistochemical detection of EYFP in liver samples from R26-lsl-EYFP reporter mice 4 days after injection of 10^8^, 10^9^ (low titer), 10^10^, 10^11^ (high titer) or 10^12^ AAV8-TBG-CRE particles. Bar: 100 µm. **(B)** Kaplan Meyer disease-free survival curves for mice of the indicated genotypes injected i.v. with low-titer AAV8-TBG-CRE at 6-8 weeks of age. The number of mice (n) and the median survival are indicated. p-values were calculated with the log-rank test. **(C)** Hematoxylin and Eosin staining of representative liver sections from tumor-bearing mice of the indicated genotypes. Bars: 400 µm (H&E 5x) or 100 µm (H&E 5x). Each tumor is identified by its unique reference number.

**Supplementary Figure 4.**
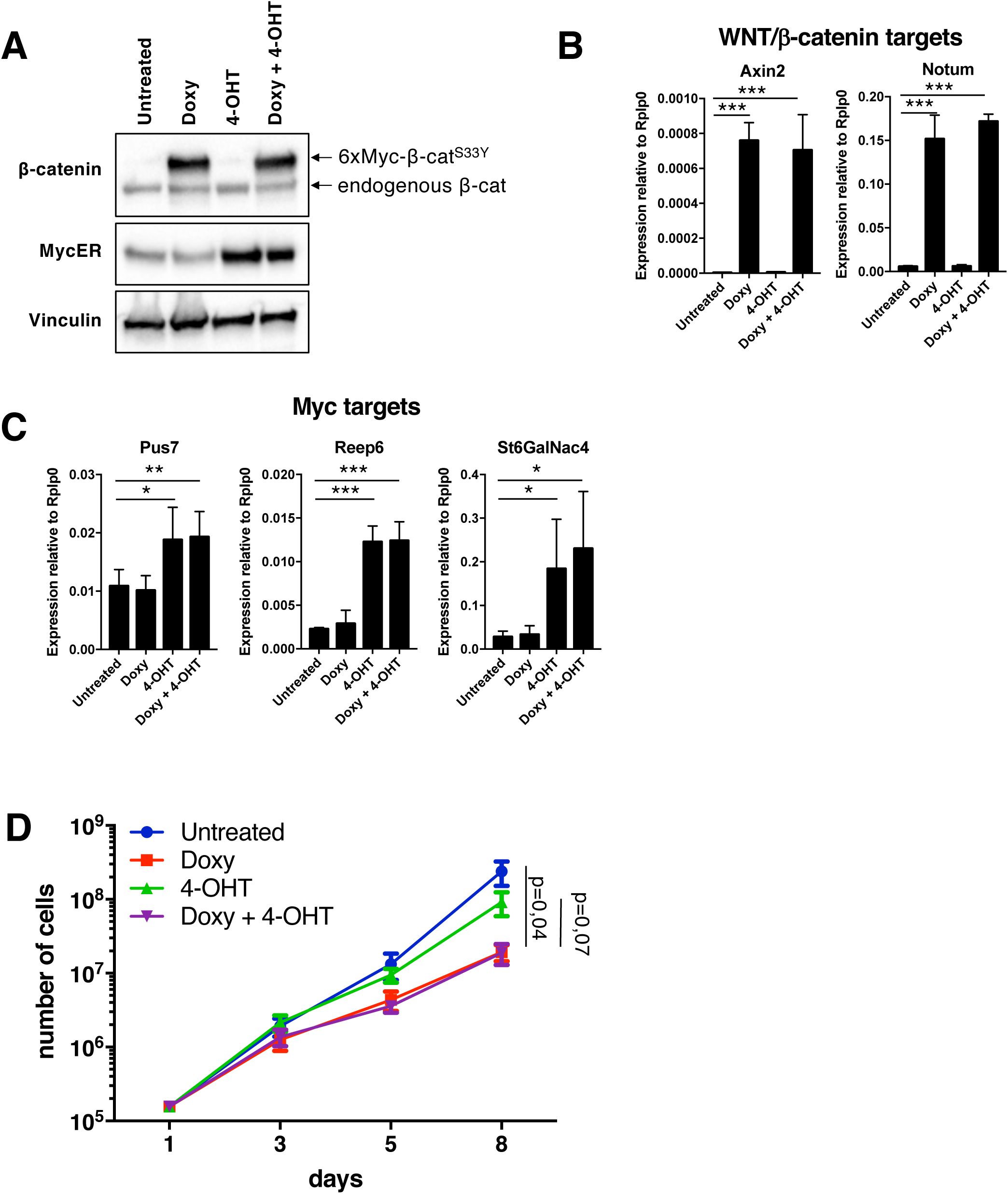
MycER and β-cat^S33Y^ activation in fibroblasts does not affect adherent-cell growth *in vitro*. **(A)** Western blot analysis of β-catenin and MycER expression in 3T9^MycER;S33Y^ cells treated for 24 hours with 1 μM Doxycycline to activate β-cat^S33Y^ and/or for 4 hours with 400nM 4-OHT to activate MycER. Note that β-cat^S33Y^ is fused to a 6xMyc-tag, explaining its slower migration relative to endogenous β-catenin. Vinculin was used as loading control. **(B-C)** RT-qPCR analysis of selected WNT/β-catenin and MYC-activated genes, as indicated, in samples from the same cells as in **(A)** Average and standard deviation for at least 3 biological replicates. **(D)** Growth curves for 3T9^MycER;S33Y^ cells cultivated in the presence of 1 μM Doxycycline and/or 400 nM 4-OHT. The number of cells at each time point were determined with quantitative flow cytometry in a MACS Quant Analyzer.

**Supplementary Figure 5.**
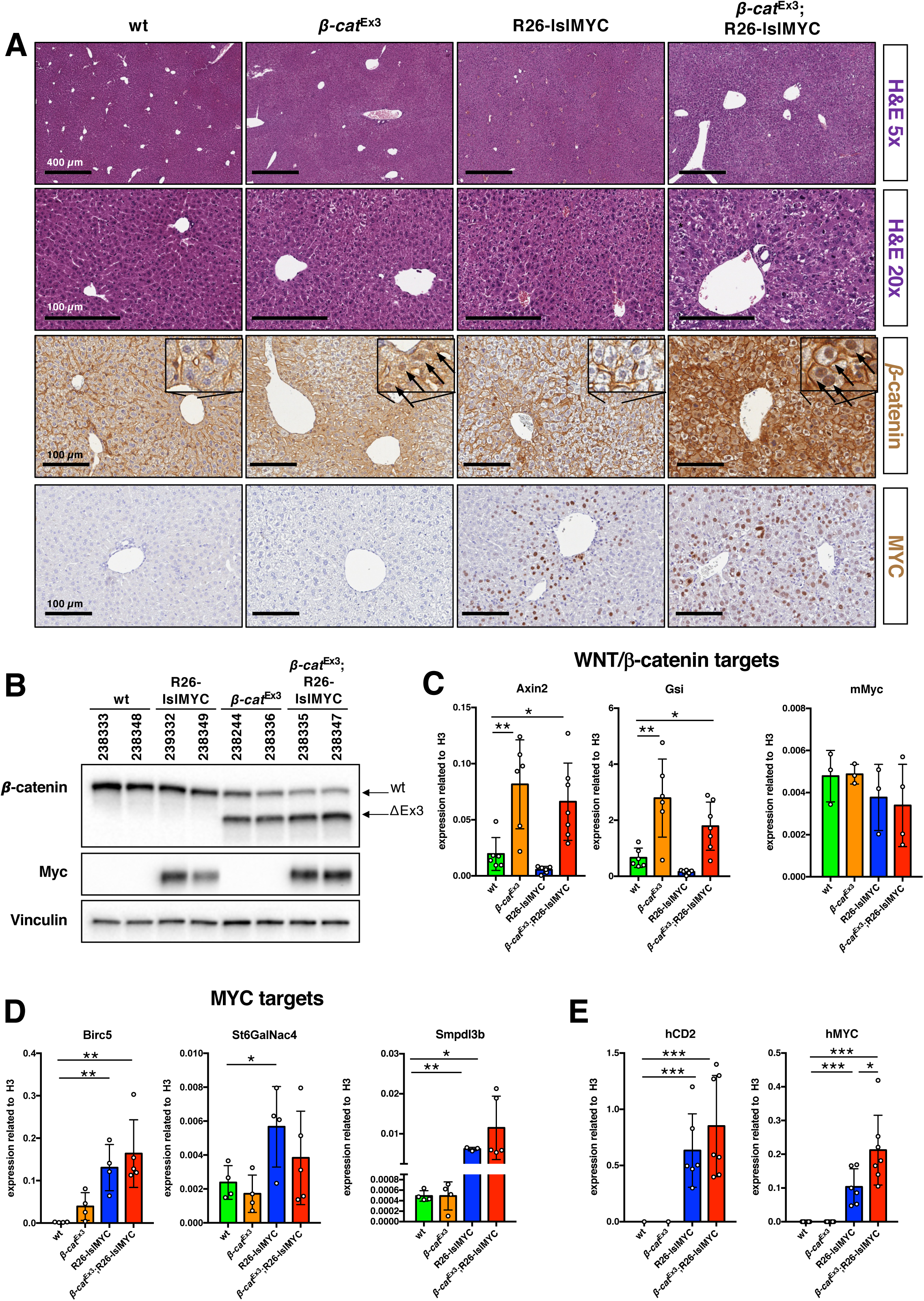
Efficient activation of MYC and β-cat^Ex3^ in the liver upon delivery of high-titer AAV8-TBG-CRE particles. **(A)** Hematoxylin and Eosin staining and immunohistochemistry for MYC and β-catenin in representative liver samples from mice of the indicate genotypes, injected with high-titer AAV8-TBG-CRE particles 4 days prior to collection. The arrow point to examples of β-catenin nuclear staining. Bars: 400 µm (H&E 5x) or 100 µm (H&E 5x, β-catenin and MYC). **(B)** Western blot analysis of β-catenin and MYC protein expression in liver lysates from mice of the indicated genotypes, treated as in **(A)** Vinculin was used as loading control. Each sample is identified by its unique reference number. **(C-E)** RT-qPCR analysis of selected mRNAs in livers of the indicated genotypes, with wild-type liver as control: **(C)** representative WNT/β-catenin-activated genes; **(D)** Representative MYC-activated genes; **(E)** human MYC and CD2, encoded by the R26-lslMYC allele (see Supplementary Figure 1A). The bar plot represents the averages and standard deviations of at least 3 biological replicates.

**Supplementary Figure 6.**
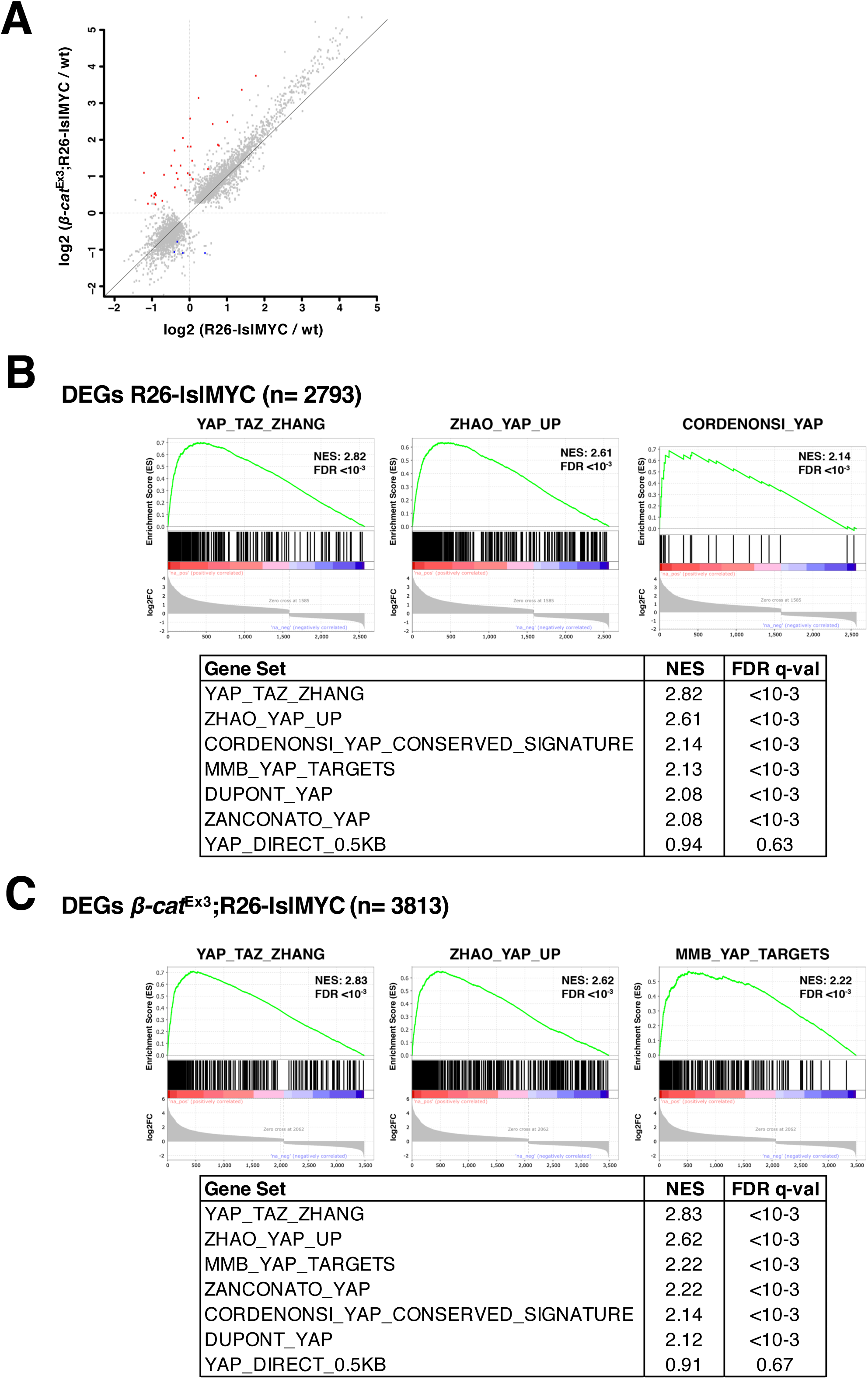
Short-term overexpression of MYC, either alone or with β-cat^Ex3^, elicits a Yap/Taz transcriptional program in the liver. **(A)** Scatter plot showing the variation in gene expression (log2FC) in R26-lslMYC (x-axis) and *β-cat*^Ex3^;R26-lslMYC (y-axis) relative to wt mice (y-axis). Genes called as DEG (qval<0.05) in either sample are shown in grey. The red and blue dots show the genes that score as up- and down-regulated, respectively, in the direct comparison between R26-lslMYC and *β-cat*^Ex3^;R26-lslMYC. **(B-C)** Gene set enrichment plots for the top 3 examples and complete list of NES and FDR values for the 7 manually curated gene lists of Yap/Taz targets. Genes scoring as DEGs (qval<0.05) in R26-lslMYC **(B)** or *β-cat*^Ex3^;R26-lslMYC **(C)** relative to the wild-type control were sorted from left to right according to the log2FC in their expression.

**Supplementary Figure 7.**
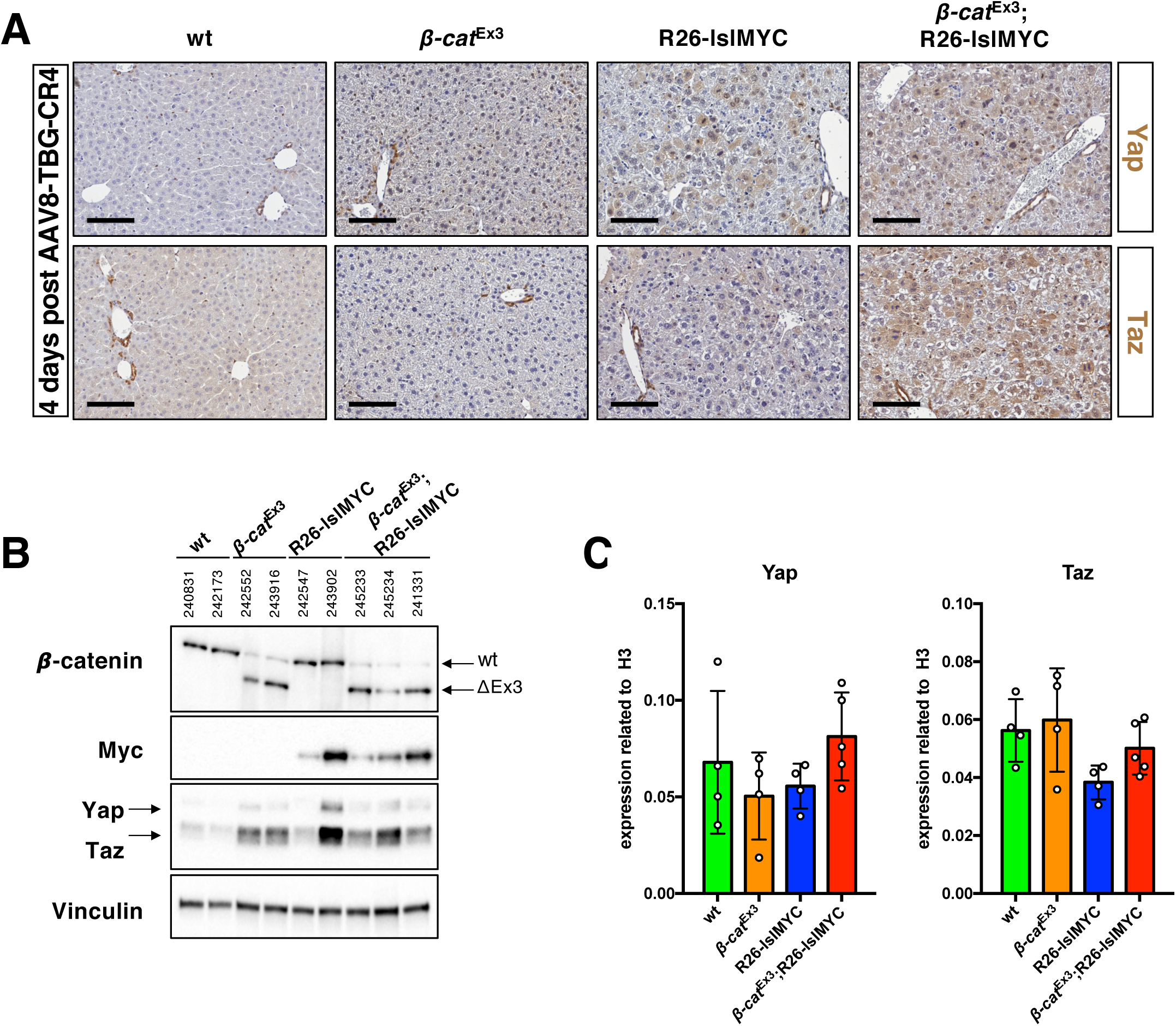
Increased Yap/Taz expression upon MYC and β-cat^Ex3^ activation in hepatocytes. **(A)** Immunohistochemical detection of Yap and Taz in representative liver samples from mice of the indicate genotypes, injected with high-titer AAV8-TBG-CRE 4 days prior to collection. Bar: 100 µm. **(B)** Western blot analysis of Yap, Taz, β-catenin and MYC protein expression in lysates from mice treated as in (A) Vinculin was used as loading control. Each sample is identified by its unique reference number. **(C)** RT-qPCR analysis of the Yap and Taz mRNAs in the same samples. The bar plots represent average and standard deviation from at least 3 biological replicates.

**Supplementary Figure 8.**
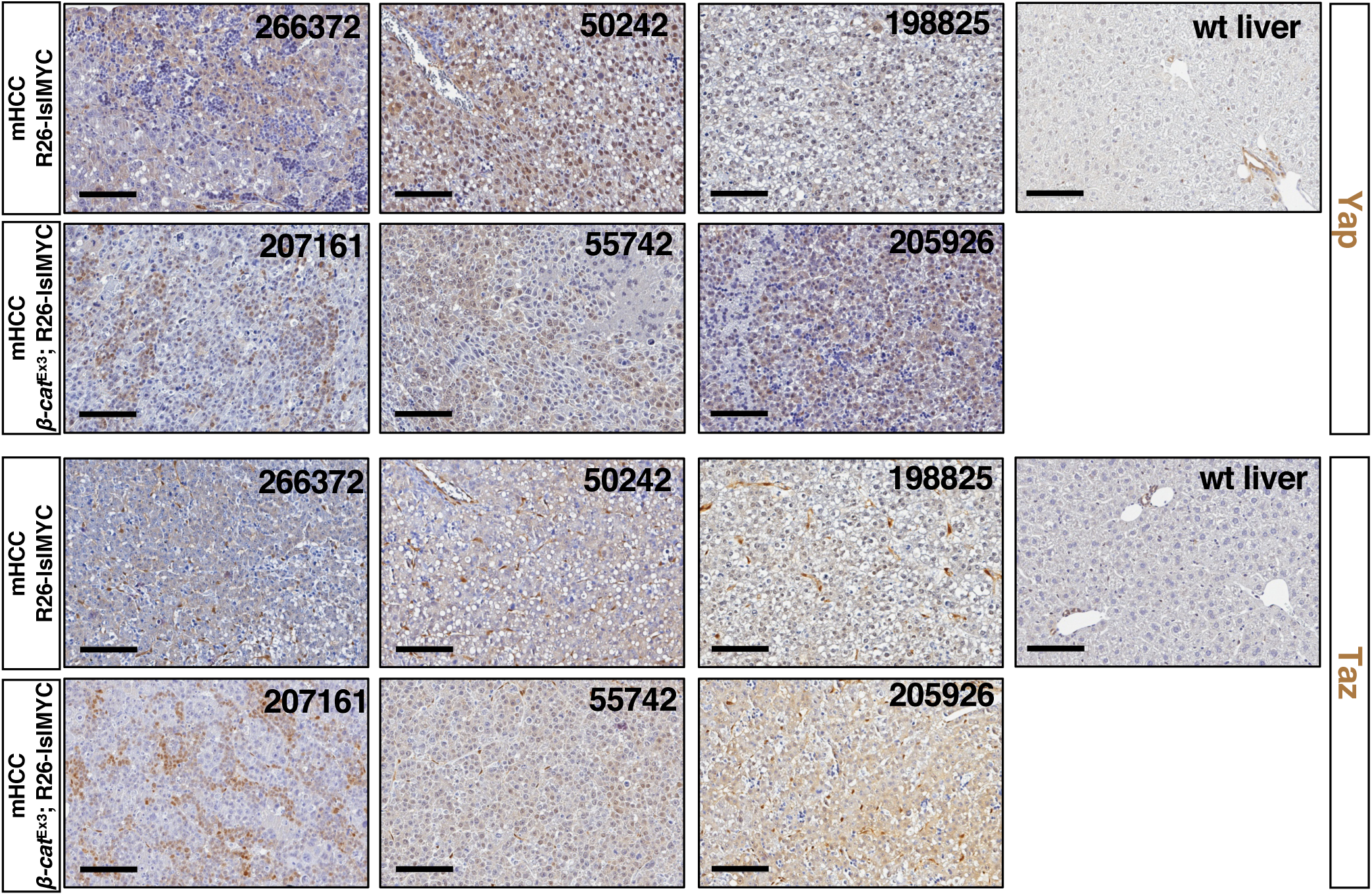
Increased Yap/Taz expression in MYC and β-cat^Ex3^-induced tumors. Immunohistochemical detection of Yap and Taz in representative mHCCs of the indicated genotypes (all in the presence of the Alb-CreER^T2^ transgene). Each tumor is identified by its unique reference number. Bar: 100 µm.

**Supplementary Figure 9.**
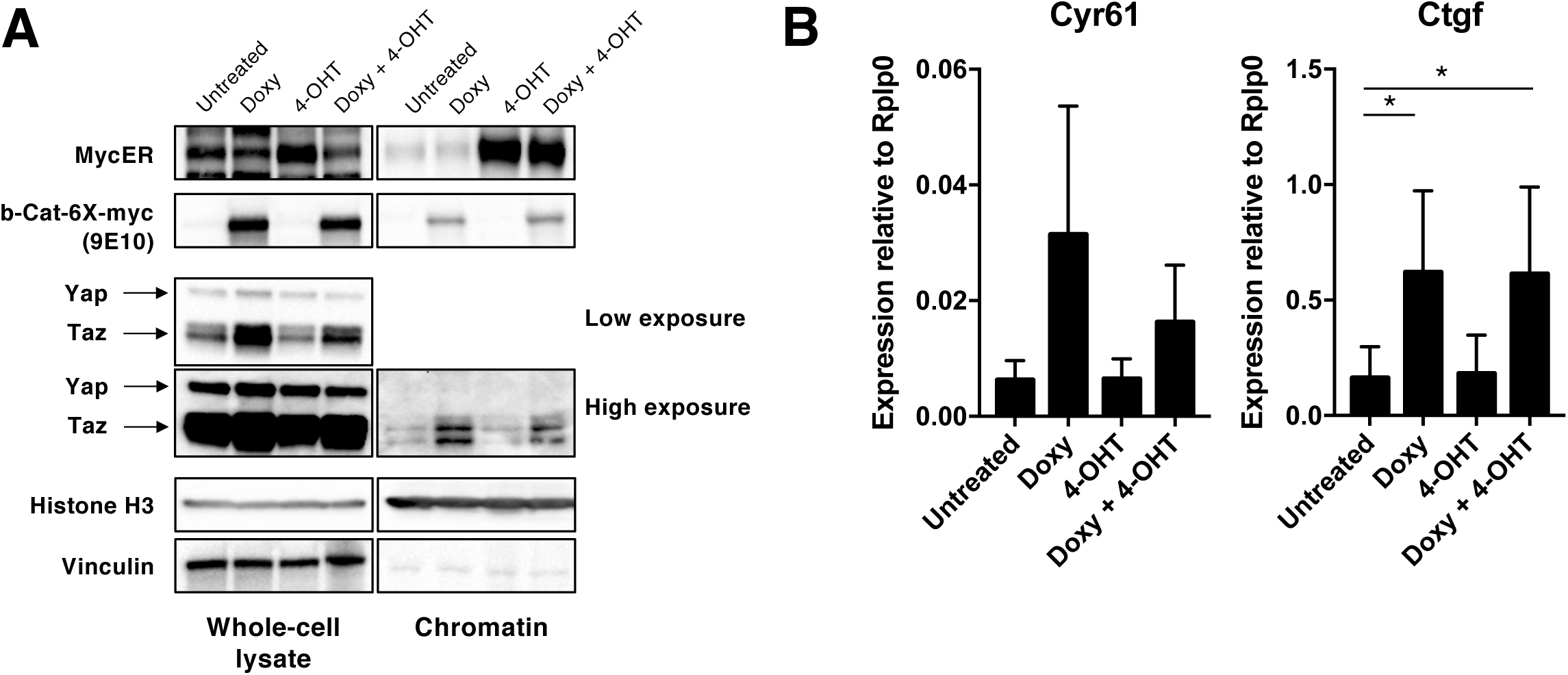
Yap/Taz are activated upon β-cat^S33Y^ overexpression in fibroblasts. **(A)** Western blot analysis of β-catenin, MycER, Yap and Taz in whole-cell lysates or chromatin associated fractions of 3T9^MycER;S33Y^ cells. The cells were treated with 1 μM Doxycycline for 24 hours to activate β-cat^S33Y^ and/or 400nM 4-OHT for 4 hours to activate MycER. Vinculin and Histone H3 were used as fractionation and loading controls. **(B)** RT-qPCR analysis of two canonical Yap/Taz transcriptional targets in the same cells. The bar plot represents averages and standard deviations for at least 3 biological replicates.

**Supplementary Figure 10.**
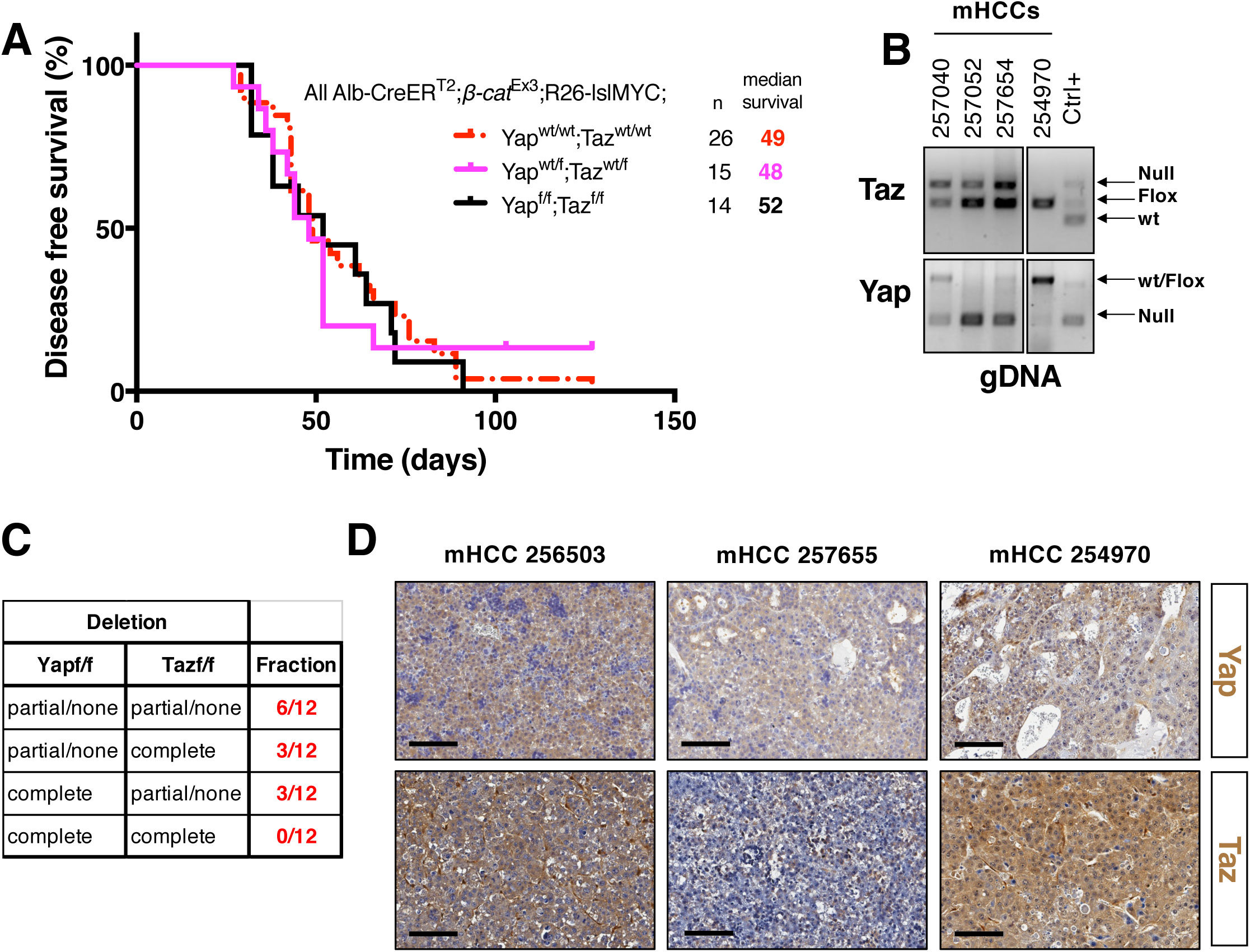
Yap/Taz are required for MYC/β-cat^Ex3^-dependent liver tumorigenesis. **(A)** Kaplan Meyer disease-free survival curves for mice of the indicated genotypes (all in the presence of the Alb-CreER^T2^ transgene). The numbers of mice (n) and the median survival are indicated. p-values were calculated with the log-rank test. **(B)** Representative results from the analysis of Cre-mediated recombination of the Yap^f/f^ and Taz^f/f^ alleles, performed by semi-quantitative PCR on genomic DNA extracted from bulk liver tumors (each identified by its unique number). As positive control (Ctrl+) for these PCRs we used a 1:1 mix of genomic DNA extracted from peripheral blood cells of Mx1-CRE^+^;Yap^f/wt^;Taz^f/wt^ mice 30 days after injection of 250 μg of poli I:C (leading to Yap^f^/Taz^f^ recombination) or vehicle (Yap^f^/Taz^f^ non-recombined)^21^. **(C)** Summary of the recombination analysis (performed as in B) on a total of 12 Alb-CreER^T2^;*β-cat*^Ex3^;R26-lslMYC;Yap^f/f^;Taz^f/f^ mHCCs. The table shows the fraction of mHCCs with the indicated recombination status at the targeted *Yap* and *Taz* loci. Complete: both alleles deleted; Partial: only one allele deleted; None: both alleles intact. **(D)** Immunohistochemical detection of Yap and Taz in representative tumor samples from Alb-CreER^T2^;*β-cat*^Ex3^;R26-lslMYC;Yap^f/f^;Taz^f/f^ mice. Each tumor is identified by its unique reference number. Bar: 100 µm.

**Supplementary Figure 11.**
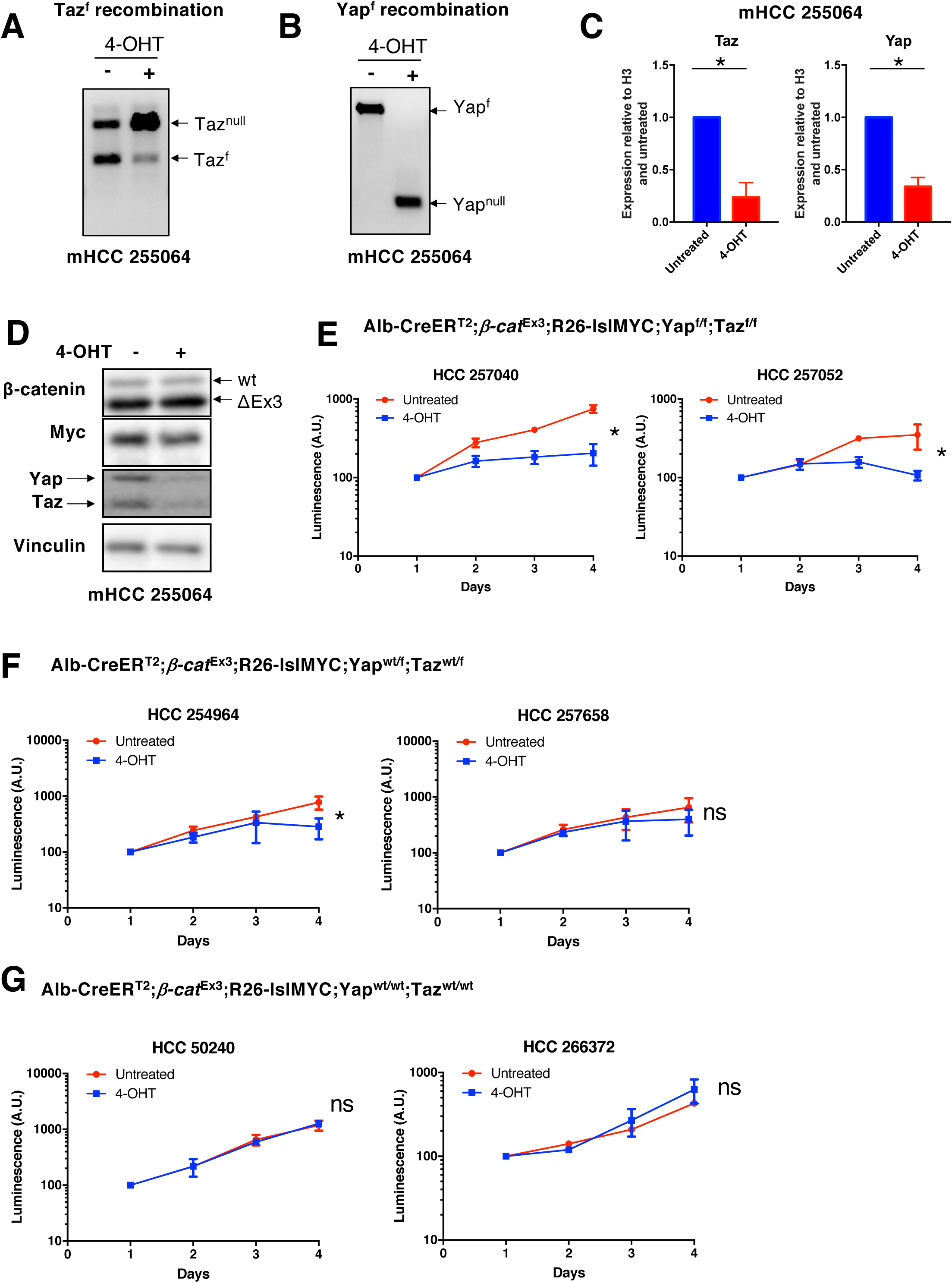
Yap/Taz are required for the survival of MYC/*β-cat*^Ex3^ mHCC tumor cells. Representative results from the analysis of Cre ER^T2^-mediated recombination of the Taz^f/f^ **(A)** or Yap^f/f^ alleles **(B)** in 4-OHT-treated Alb-CreER^T2^;*β-cat*^Ex3^;R26-lslMYC;Yap^f/f^;Taz^f/f^ tumor cells (mHCC 255064). When indicated, cultures were treated for 24h with 4-OHT in order to acutely activate CreER^T2^. The status of the alleles was analyzed by semi-quantitative PCR on genomic DNA. **(C)** RT-qPCR analysis of the expression of *Yap* and *Taz* mRNA levels in the same cells. **(D)** Representative Western blot analysis of Yap, Taz, β-catenin and MYC expression in lysates from mHCC 255064 (Alb-CreER^T2^;*β-cat*^Ex3^;R26-lslMYC;Yap^f/f^;Taz^f/f^) treated or not for 24h with 400nM 4-OHT, as indicated. Vinculin was used as loading control. **(E-G)** Growth curves for two independent mHCC cultures of each genotype: **(E)** Alb-CreER^T2^;*β-cat*^Ex3^;R26-lslMYC;Yap^f/f^;Taz^f/f^; **(F)** Alb-CreER^T2^;*β-cat*^Ex3^;R26-lslMYC;Yap^wt/f^;Taz^wt/f^; (**G)** Alb-CreER^T2^;*β-cat*^Ex3^;R26-lslMYC;Yap^wt/wt^;Taz^wt/wt^. All cells were cultured in the absence (red line) or presence (blue line) of 400 nM 4-OHT to activate the CreER^T2^ recombinase. Relative cell numbers were determined with the Cell Titer Glo assay. A.U.: Arbitrary Units.

**Supplementary Figure 12.**
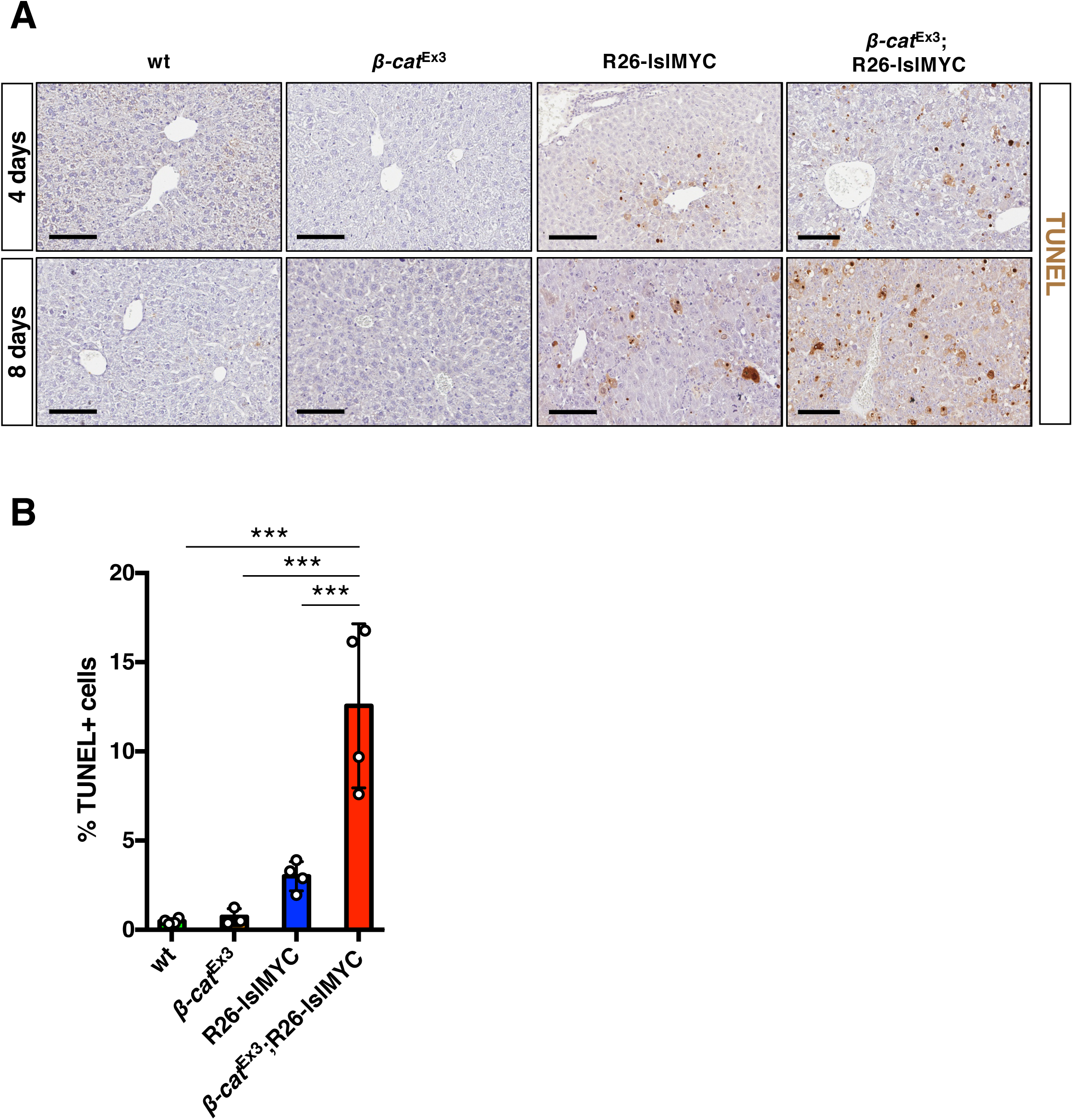
β-cat^Ex3^ activation does not dampen MYC-dependent apoptosis in hepatocytes. **(A)** Representative images of TUNEL staining for apoptotic cells in liver samples from mice of the indicated genotypes, injected with AAV8-TBG-CRE either 4 or 8 days prior to collection. Bar: 100 µm. **(B)** Quantification of TUNEL positive cells 8 days post AAV8-TBG-CRE injection the same animals as in (A). The bar plot represents average and standard deviation from 3 biological replicates.

## Notes

**<u>Financial Support</u>:** This study was supported by funding from the European Research Council (grant agreement no. 268671-MYCNEXT), the Italian Health Ministry (RF-2011-02346976) and the Italian Association for Cancer Research (AIRC, IG 2012-13182 and 2015-16768) to B.A., from Worldwide Cancer Research (15-1260) to A.S. and from AIRC to S.C. (IG 2018-21663).

